# Tyrosine phosphorylation regulates hnRNPA2 granule protein partitioning & reduces neurodegeneration

**DOI:** 10.1101/2020.03.15.992768

**Authors:** Veronica H. Ryan, Theodora Myrto Perdikari, Mandar T. Naik, Camillo F. Saueressig, Jeremy Lins, Gregory L. Dignon, Jeetain Mittal, Anne C. Hart, Nicolas L. Fawzi

## Abstract

mRNA transport in neurons is a ubiquitous process but has been often overlooked as a contributor to disease. Mutations of transport granule protein hnRNPA2 cause hereditary proteinopathy of neurons, myocytes, and bone. Here, we examine transport granule component specificity, assembly/disassembly, and the link to neurodegeneration. hnRNPA2 transport granule components hnRNPF and ch-TOG interact weakly with hnRNPA2 yet they each partition specifically into hnRNPA2 liquid phases. hnRNPA2 tyrosine phosphorylation dissociates granule interactions by reducing hnRNPA2 phase separation and preventing partitioning of hnRNPF and ch-TOG; tyrosine phosphorylation also decreases aggregation of hnRNPA2 disease mutants. A *C. elegans* model of hnRNPA2 D290V-associated neurodegeneration exhibits TDP-43 ortholog-dependent glutamatergic neurodegeneration. Expression of the tyrosine kinase that phosphorylates hnRNPA2 reduces glutamatergic neurodegeneration. The evidence for specific partitioning of granule components as well as disruption of these interactions and reduction of neurodegeneration by tyrosine phosphorylation suggest transport granule biology has a role in the pathogenesis of neurodegeneration.

## Introduction

Cells, particularly highly polarized cells like neurons, organize their cytosol using both membrane-bound organelles and membraneless organelles (MLOs), condensates of RNA and proteins (Banani et al., 2017). Neurodegenerative diseases including amyotrophic lateral sclerosis and frontotemporal dementia (ALS/FTD) have been linked to disruption of the components and properties of MLOs, possibly through MLO stabilization by mutations in proteins capable of liquid-liquid phase separation (LLPS), a phenomenon where proteins and nucleic acids demix from the surrounding solution (Nedelsky and Taylor, 2019; Ryan and Fawzi, 2019). The relationship between stress granules, MLOs observed in cells after exposure to exogenous stress, and these these ALS/FTD-associated proteins, including FUS, TDP-43, hnRNPA1, and hnRNPA2, has been the primary focus of work relating MLOs to neurodegeneration (Burke et al., 2015; Conicella et al., 2016; Molliex et al., 2015; Monahan et al., 2017; Patel et al., 2015; Ryan et al., 2018; Wang et al., 2018a). However, many of these disease-associated proteins are also found in physiological MLOs observed in the absence of stress, notably transport granules.

Transport granules, called neuronal granules when found in neurons, move mRNAs from the perinuclear space to sites of local translation. Local translation is important in many cell types but is critical for myelination by oligodendrocytes and neuronal functions, including synaptic plasticity and axon guidance. Different kinds of mRNA transport granules likely exist, with distinct mRNA cargos, including neuronal β-actin mRNA transport granules containing zip-code binding protein/IGF2BP1 (Elvira et al., 2006; Kiebler and Bassell, 2006; Zhang et al., 2001) and hnRNPA2-containing granules transporting myelin basic protein mRNA in oligodendrocytes or Arc, neurogranin, and αCamKII in neurons (Ainger et al., 1993; Brumwell et al., 2002; Gao et al., 2008; Shan et al., 2003). Several other protein components of hnRNPA2-containing transport granules have been identified, including hnRNPF, hnRNPAB, hnRNPE1, hnRNPK, and the microtubule associated protein ch-TOG (CKAP5) (Francone et al., 2007; Kosturko et al., 2005; Kosturko et al., 2006; Laursen et al., 2011; Raju et al., 2011; Torvund-Jensen et al., 2014; White et al., 2012). Importantly, hnRNPA2 transport granules appear to contain a specific set of proteins and exclude related proteins associated with other cytoplasmic granules (e.g. other RNA-binding proteins found in stress granules), yet the mechanisms of granule component specificity remain unclear. Some of these protein interactions with hnRNPA2 are RNA-dependent (Laursen et al., 2011; Raju et al., 2011; Torvund-Jensen et al., 2014) but hnRNPF and ch-TOG both directly bind to hnRNPA2 (Falkenberg et al., 2017; Kosturko et al., 2005; White et al., 2012). Interactions within the granules are regulated; mRNA is released for local translation in processes when hnRNPA2 and hnRNPF are locally phosphorylated by the tyrosine kinase Fyn (White et al., 2012; White et al., 2008). Yet, the molecular basis for both hnRNPA2-hnRNPF and hnRNPA2-TOG interactions and their disruption by phosphorylation remains unknown.

hnRNPA2 mutations cause multisystem proteinopathy (MSP), a degenerative disease where patients experience ALS/FTD, with inclusion body myopathy, and Paget’s disease of bone (PDB) (mutation: D290V) (Kim et al., 2013) as well as PDB alone in a separate family (mutation: P298L) (Qi et al., 2017). These disease mutations drive aggregation of the protein *in vitro* (Kim et al., 2013; Ryan et al., 2018) and D290V alters the association of hnRNPA2 with stress granules (Kim et al., 2013). hnRNPA2 contains two RNA recognition motifs (RRMs) and a glycine-rich low complexity (LC) domain, which is necessary and sufficient for LLPS (Ryan et al., 2018). Importantly, both mutations are located in this aggregation-prone “prion-like” LC domain (named for its enrichment in polar residues found in yeast prion proteins) (King et al., 2012). Given the ability of hnRNPA2 to undergo LLPS, we set out to determine the molecular basis for interactions of hnRNPA2 with hnRNPF and TOG by evaluating their ability to specifically co-phase separate into *in vitro* models of reconstituted multi-component hnRNPA2 transport granules. Furthermore, as posttranslational modifications (PTMs) alter phase separation and aggregation (Monahan et al., 2017; Nott et al., 2015; Ryan et al., 2018; Wang et al., 2018a), we tested the hypothesis that tyrosine phosphorylation disrupts the interactions between granule components, leading to granule dissociation, and prevents toxic aggregation of the disease mutants *in vitro* and *in vivo*. We also hypothesized that these interactions would alter neurodegeneration in an animal model. Here, we used nuclear magnetic resonance (NMR) spectroscopy, molecular simulation, *in vitro* phase separation assays, and a novel *C. elegans* model to probe the molecular basis for hnRNPA2-containing transport granule assembly, disassembly, and hnRNPA2-associated neurodegeneration. These studies point to the possibility for modulation of transport granule component interactions as a therapeutic strategy in neurodegenerative disease.

## Results

### hnRNPA2 arginine residues are required for the interaction with transport granule component hnRNPF

hnRNPA2 was previously shown to undergo LLPS (Ryan et al., 2018) and interact with other protein components of myelin basic protein mRNA transport granules, including hnRNPF (White et al., 2012). Importantly, hnRNPF interacts directly with hnRNPA2 protein in transport granules (White et al., 2012), yet the biophysical details of this interaction between two these prion-like domain containing proteins remain unclear. Here we sought to reconstitute and structurally characterize this interaction.

First, we tested which domains mediate the hnRNPA2-hnRNPF interaction. Using purified recombinant hnRNPA2 low complexity domain (LC, residues 190-341) and full length (FL) with hnRNPF prion-like domain (PLD, resides 365-415), and FL, we asked if fluorescently tagged hnRNPF partitions into hnRNPA2 droplets. hnRNPF (PLD or FL) did not undergo LLPS alone in the conditions and concentrations required for hnRNPA2 LC or FL LLPS (Figure 1A). However, hnRNPF PLD partitioned into hnRNPA2 LC (Figure 1A) and FL (Figure S1A) droplets and was equally distributed throughout the droplets. hnRNPF FL partitioned into both hnRNPA2 LC and hnRNPA2 FL droplets (Figure S1A-B). The PLD of hnRNPF was not essential for partitioning into hnRNPA2 droplets, as hnRNPF lacking the PLD (hnRNPF ΔPLD, residues 1-364) partitioned into hnRNPA2 FL droplets (Figure S1A). We attempted to test if the hnRNPF PLD is necessary for partitioning into hnRNPA2 LC droplets and while some co-localization was observed (Figure S1A), hnRNPF ΔPLD seemed to aggregate, possibly because the hnRNPA2 LC LLPS assay conditions required crossing the hnRNPF ΔPLD predicted isoelectric point (pI), where proteins are likely to self-assemble. Taken together, these data suggest that the PLD of hnRNPF avidly partitions into hnRNPA2 droplets.

**Figure 1:**
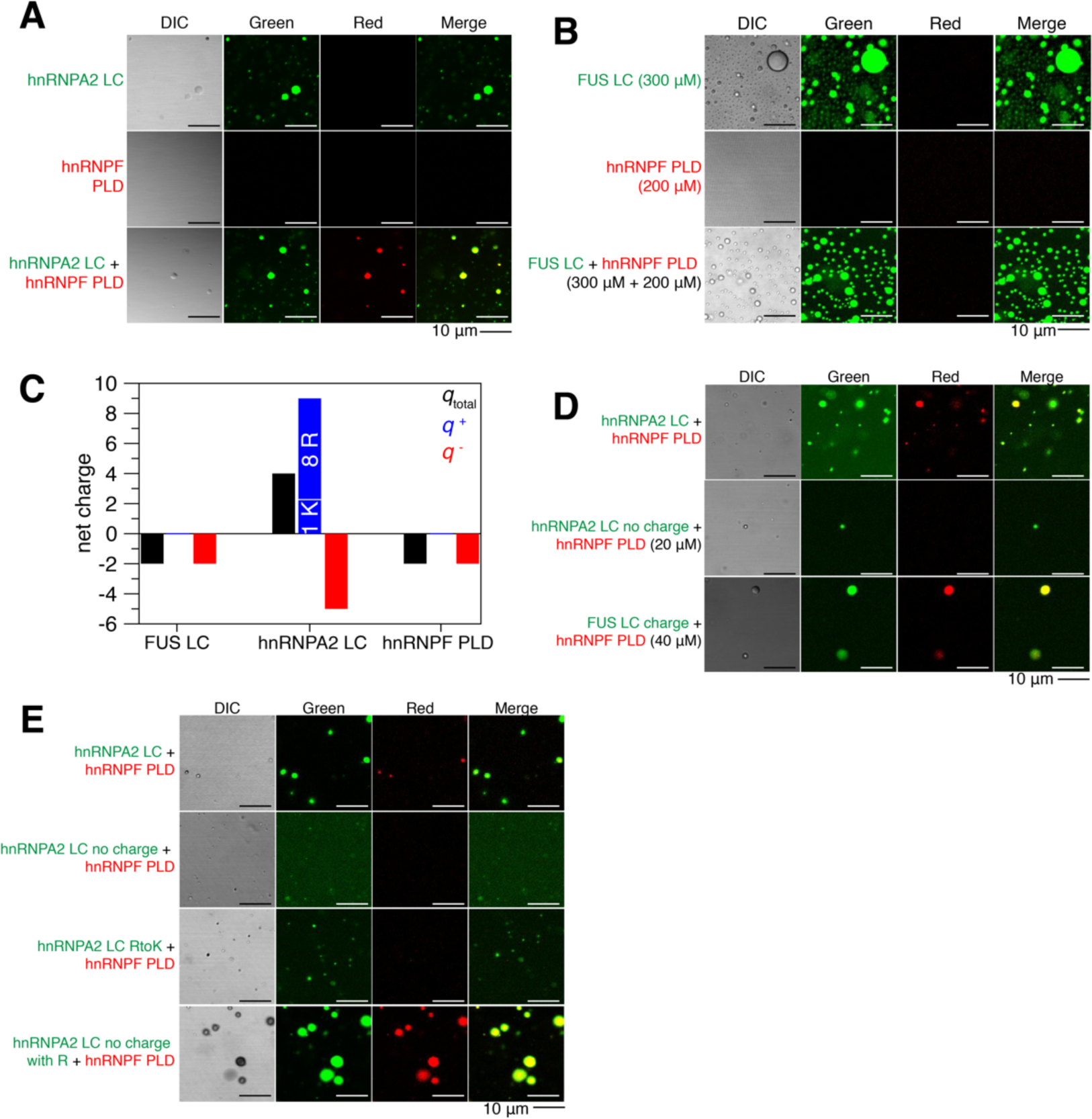
Arginine in hnRNPA2 LC is required for the interaction with granule component hnRNPF PLD. See also Figure S1. (A) hnRNPA2 LC (AlexaFluor488-tagged, green) undergoes LLPS, while hnRNPF PLD (AlexaFluor555-tagged, red) does not. However, hnRNPF PLD partitions into hnRNPA2 LC droplets when mixed at a 1:1 ratio. Conditions: 20 µM indicated protein (∼1% fluorescently tagged), 20 mM MES pH 5.5, 50 mM NaCl, 150 mM urea. Scale bar: 10 µm. (B) At 300 µM, FUS LC (AlexaFluor488-tagged, green) undergoes LLPS, but at 200 µM hnRNPF PLD (AlexaFluor555-tagged, red) still does not undergo LLPS. When mixed at 300 µM FUS LC and 200 µM hnRNPF PLD, hnRNPF PLD does not partition into FUS LC droplets. Conditions: 300 µM FUS and 200 µM hnRNPF PLD (∼1% fluorescently tagged), 20 mM MES pH 5.5, 150 mM NaCl, 150 mM urea. Scale bar: 10 µm. (C) While FUS LC and hnRNPF PLD both have a small negative predicted net charge at physiological pH, hnRNPA2 LC has a predicted +4 net positive charge, due to the 9 positively charged residues (8 arginine, 1 lysine) and 5 negatively charged residues. (D) Removal of the charged residues from hnRNPA2 LC (hnRNPA2 LC no charge) prevents partitioning of hnRNPF PLD into the hnRNPA2 LC phase. Addition of hnRNPA2 LC-like charged residue patterning to FUS LC allows the partitioning of hnRNPF PLD at 40 µM. Conditions: 20 µM hnRNPA2 LC and hnRNPA2 LC no charge, 40 µM FUS LC charge, hnRNPF PLD concentration matches other protein in mixture (either 20 or 40 µM) (all ∼1% fluorescently tagged), 20 mM MES pH 5.5 50 mM NaCl, 150 mM urea. Scale bar: 10 µm. (E) Substitution of all arginines in hnRNPA2 LC with lysine prevents the partitioning of hnRNPF PLD into hnRNPA2 LC RtoK droplets. Removing all charged residues except for arginine from hnRNPA2 LC (A2 LC no charge with R) allows partitioning of hnRNPF PLD into droplets, indicating arginine in hnRNPA2 LC is required and necessary for hnRNPF partitioning. hnRNPA2 LC RtoK does not phase separate much as hnRNPA2 LC at these conditions, see **Figure S1J** for quantification of phase separation of variants. Conditions: 20 µM proteins, 20 mM MES pH 5.5 50 mM NaCl, 150 mM urea. Scale bar: 10 µm.

Next, we focused on delineating the molecular basis for the interaction between hnRNPA2 and hnRNPF. To visualize hnRNPF PLD with residue-by-residue resolution, we performed multidimensional NMR spectroscopy to observe each backbone amide position. From the narrow chemical shift dispersion and secondary shifts, we conclude that hnRNPF PLD is primarily disordered (Figure S1C-D). Then, we attempted to localize the sites of interactions between hnRNPA2 LC and hnRNPF PLD using NMR titrations. At low salt conditions where hnRNPA2 LC alone does not phase separate, addition of hnRNPF PLD induced LLPS of hnRNPA2 LC (Figure S1E), reminiscent of how Fyn-SH3 a folded domain that does not phase separate on its own, also induces LLPS of hnRNPA2 LC (Amaya et al., 2018). The titrations showed a distributed interaction across the entire length of hnRNPF PLD and hnRNPA2 LC and did not implicate specific residues or residue types in hnRNPA2 LC interactions with hnRNPF PLD (Figure S1E-F), consistent with the repetitive, low complexity sequence of these PLDs.

An important question in the field of MLOs is the origin of specificity of granule partitioning. Although no specific sites of interaction were observed by NMR, we wondered if the sequences of hnRNPA2 LC and hnRNPF PLD encode any specificity in partitioning. Therefore, we examined specificity by testing if hnRNPF PLD could co-phase separate with the FUS LC. FUS is also found in stress granules but not known to be in hnRNPA2-myelin basic protein mRNA granules. Importantly, FUS LC has different amino acid composition than that of hnRNAP2. However, we found that FUS LC and FUS LC 12E, a phosphomimetic form of FUS that does not undergo LLPS (Monahan et al., 2017), are both capable of partitioning into hnRNPA2 (Figure S1G). Interestingly, when hnRNPF PLD was mixed with FUS LC, it did not partition into FUS LC droplets (Figure 1B), demonstrating specificity of the hnRNPA2-hnRNPF partitioning. We hypothesized that charged residues might underly this specificity, as both FUS LC and hnRNPF PLD are slightly negatively charged, while hnRNPA2 LC has a net positive charge (Figure 1C). To test the role of charged residues in hnRNPA2 LC specifying the interaction with hnRNPF, we changed almost all the charged residues from hnRNAP2 LC to serine or glutamine resulting in a “FUS LC-like” charge (which we termed hnRNPA2 LC no charge) or changed serine/glutamine residues in FUS to charged residues to give it “hnRNPA2 LC-like” pattern of charged residues (FUS LC charge) and then examined hnRNPF PLD partitioning into these charged residue variants. hnRNPF PLD did not partition into hnRNPA2 LC no charge but did partition into FUS LC charge (Figure 1D, S1H), consistent with our hypothesis.

Given that arginine contacts with aromatic residues are important for phase separation in hnRNPA2 (Ryan et al., 2018) and other proteins (Vernon et al., 2018; Wang et al., 2018b), we hypothesized that partitioning of hnRNPF PLD into hnRNPA2 LC droplets required arginine in hnRNPA2 LC and tyrosine in hnRNPF PLD. Therefore, we changed hnRNPF PLD tyrosines to serine (YtoS), hnRNPA2 LC arginines to lysine (RtoK), and also tested a form of hnRNPA2 LC with arginine residues retained but otherwise “FUS LC-like” depletion of charged residues (hnRNPA2 LC no charge with R). First, we found that hnRNPF PLD YtoS still partitioned into hnRNPA2 LC droplets, although possibly more weakly than hnRNPF PLD WT (Figure 1E, S1I), implying that hnRNPF PLD tyrosines may be important for partitioning but are not the only residue type contributing to the interaction between hnRNPA2 LC and hnRNPF PLD. Second, we found that replacing arginine with lysine in hnRNPA2 LC to maintain net charge and charge patterning (RtoK) prevented partitioning of hnRNPF PLD or hnRNPF PLD YtoS, suggesting that arginine-specific contacts and not just positive charge determine partitioning. Third, we found that adding back arginine to hnRNPA2 LC no charge allowed partitioning of hnRNPF PLD and hnRNPF PLD YtoS (Figure 1E). Inconveniently, FUS LC charge RtoK or FUS LC with R did not undergo LLPS at the same conditions as hnRNPA2 LC, so we could not perform the complementary experiments testing hnRNPF PLD partitioning into modified FUS constructs (Figure S1I). We also tested the role of asparagine residues in hnRNPA2 LC by changing asparagine to serine (hnRNPA2 LC NtoS) and the effect of serine residues in hnRNPF PLD by changing serine to alanine (StoA), but these changes did not alter partitioning (Figure S1I). We conclude that arginine residues in hnRNPA2 LC are critical to specify co-phase separation of hnRNPF PLD.

### hnRNPA2 LC interacts with TOG D1 weakly through its disordered loops and helical face

Similar to hnRNPF, ch-TOG was previously shown to interact with hnRNPA2 in myelin basic protein transport granules (Falkenberg et al., 2017; Francone et al., 2007; Kosturko et al., 2005). Previous work reported that hnRNAP2 interacts with each individual domain of ch-TOG at nanomolar affinity via the hnRNPA2 LC (Falkenberg et al., 2017). Given our results with hnRNPF, we hypothesized that the first domain of ch-TOG, TOG D1 (residues 1-250), would also partition into hnRNPA2 droplets. As expected for a single globular domain, recombinant TOG D1 did not undergo LLPS in the micromolar concentration range or buffers tested, but fluorescently tagged TOG D1 partitioned into hnRNPA2 LC and FL droplets (Figure 2A, S2A). To map the interactions between hnRNPA2 LC and TOG D1, we performed assignment experiments on TOG D1 and confirmed its primarily *α*-helical secondary structure (Figure S2D-E). We performed NMR titrations with hnRNPA2 LC and TOG D1 and found no detectable interaction between hnRNPA2 LC and TOG D1 in the monomeric state using this technique (Figure S2G-H). The lack of evidence for binding suggests that the interaction between these two proteins is likely in the micromolar to millimolar range, weaker than previously reported (which we attribute to artifacts due to LLPS, see discussion).

**Figure 2:**
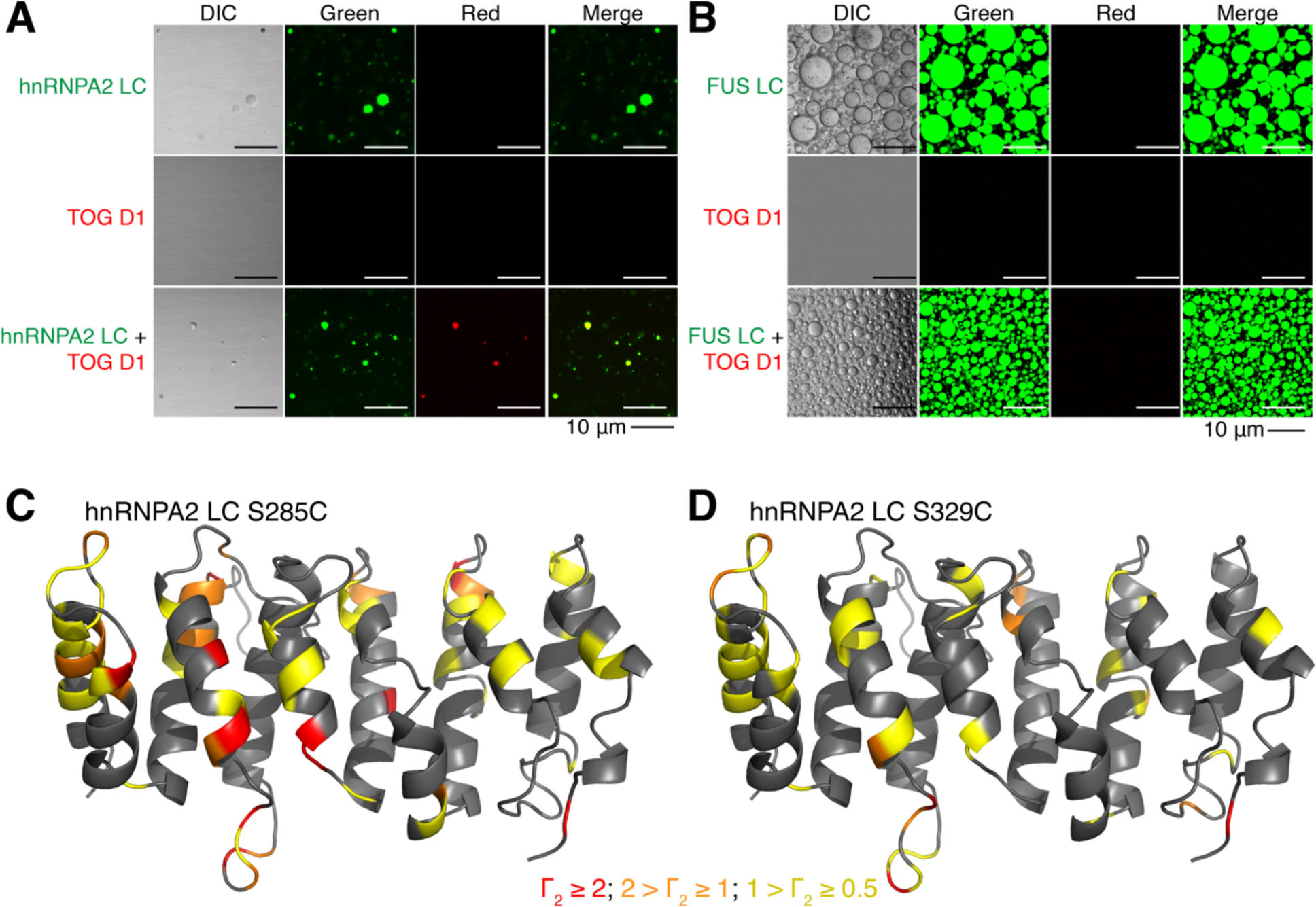
Transport granule component TOG D1 interacts weakly with hnRNPA2 LC. See also Figure S2. (A) AlexaFluor488-tagged (green) hnRNPA2 LC undergoes LLPS, while AlexaFluor555-tagged (red) TOG D1 does not. However, TOG D1 partitions into hnRNPA2 LC droplets when mixed at a 1:1 ratio. Conditions: 20 µM indicated protein (∼1% fluorescently tagged), 20 mM MES pH 5.5, 50 mM NaCl, 150 mM urea. Scale bar: 10 µm. hnRNPA2 LC control duplicated from Figure 2A as hnRNPF PLD and TOG D1 samples were made concurrently. (B) Similar to hnRNPF PLD, AlexaFluor555-tagged TOG D1 does not undergo LLPS at 300 µM or partition into AlexaFluor488-tagged FUS LC droplets with both proteins at 300 µM. Conditions: 300 µM proteins (∼1% fluorescently tagged), 20 mM MES pH 5.5 150 mM NaCl, 150 mM urea. Scale bar: 10 µm. (C-D) TOG D1 homology structure with Γ_2_ values from PRE experiments for hnRNPA2 LC (c) S285C and (d) S329C. Amino acids are colored based on Γ_2_ value: red corresponds to Γ_2_ > 2, orange to 2 > Γ_2_ > 1, yellow to 1 > Γ_2_ > 0.5.

We observed that the co-phase separation of between hnRNPA2 LC and TOG D1 is specific; TOG D1 partitioned into hnRNPA2 LC but not FUS LC droplets (Figure 2B). We hypothesized that because TOG D1 has a charged surface (Figure S2C), the charge variants of hnRNAP2 LC and FUS LC would also alter TOG partitioning. Indeed, TOG D1 partitioned into FUS LC charge droplets, yet it also partitioned into hnRNPA2 LC no charge droplets (Figure S2B). This result is different than what we observed for hnRNPF PLD and indicates that charged residues contribute to the specificity of partitioning, but other interactions contribute as well. To begin to elucidate these interactions, we turned to NMR techniques that provide position-specific information on weak, transient interactions. High resolution TROSY-based paramagnetic relaxation enhancement (PRE) experiments on mixtures of NMR invisible hnRNPA2 LC tagged with a small (∼120 Da) paramagnetic label at a single engineered cysteine site (either S285C or S329C) (Ryan et al., 2018) and NMR-visible (^2^H ^15^N) TOG D1 revealed weak interactions, as relaxation enhancement was seen at several residues of TOG D1 (Figure S2F). These interactions suggest that the region of hnRNPA2 LC bearing the paramagnetic tag comes in close proximity with particular parts of the TOG D1 surface. We then mapped the PREs on a homology model of TOG D1 and found that they localized to disordered loops and helix faces of TOG D1 (Figure 2C-D, Figure S2F). Upon sorting PREs > 0.5 s^-1^ by residue type, PREs are most often observed at non-polar positions (28/76 residues with PREs) and charged residues (27/76). However, non-polar amino acids are the most prevalent type in TOG D1 (70/165 assigned residues); after normalizing to the number of assignable residues of that type, residues with PREs are enriched in polar residues (22/40 assigned), suggesting that polar contacts are important for the interaction. Interestingly, PREs arising from hnRNPA2 LC S285C, labeled in the assembly-prone PLD, showed larger PREs than S329C, labeled in the glycine-rich tail, indicating a stronger interaction with this highly conserved region (Kim et al., 2013). hnRNPA2 LC self-interactions are also weaker at S329C than S285C (Ryan et al., 2018), so it is possible that hnRNPA2 LC interacts with TOG D1 using similar polar-residue contacts as hnRNPA2 LC uses for self-interactions.

### Tyrosine phosphorylation of hnRNPA2 LC alters LLPS and prevents partitioning of granule components

Because hnRNPA2 LC is a known target of tyrosine phosphorylation and tyrosine phosphorylation was previously shown to release mRNA from hnRNPA2 granules for translation (White et al., 2008), we hypothesized that tyrosine phosphorylation of hnRNPA2 LC would alter phase separation propensity, as we found previously for serine/threonine phosphorylation of FUS LC (Monahan et al., 2017). We generated and purified recombinant tyrosine phosphorylated (pY) hnRNPA2 LC with approximately 4 to 8 phosphorylated tyrosines (of 17 tyrosines total, by mass spectrometry) (Figure S3A). We found that pY hnRNPA2 LC undergoes LLPS with an inverse salt dependence compared to unphosphorylated hnRNPA2 LC. Specifically, pY hnRNPA2 LC phase separated more at low salt conditions and less at physiological and high salt conditions (Figure 3A-B). The high phase separation propensity at low salt conditions may be because the phosphorylated protein is at or near its pI. We also performed microscopy experiments with pY hnRNAP2 using a subset of conditions and always observed droplets (Figure 3A). pY is negatively charged. To determine if the altered LLPS of pY hnRNPA2 LC is a unique feature of phosphotyrosine or is due to increased negative charge, we tested if hnRNPA2 LC with phosphoserine mimics could also alter LLPS as a function of salt concentration. We generated two phosphomimic constructs, hnRNPA2 LC 5E, where 5 serines are changed to glutamate, and hnRNPA2 LC 12E, where all twelve serines are changed to glutamate (hnRNPA2 LC contains no threonine). At physiological ionic strength (150 mM NaCl) and above, we find that, compared to unmodified hnRNAP2 LC, 5E shows a modest reduction in LLPS while 12E shows substantially less LLPS (Figure S3F), consistent with introduction of negatively charged residues disrupting LLPS. Interestingly, at low salt conditions (0 mM NaCl), 5E phase separates, consistent with this variant being at its pI. To identify the interactions underlying LLPS propensity for pY and serine phosphomimic constructs, we performed NMR titrations of free amino acids (pY, Y, pS, R) into hnRNPA2 LC. Due to the high propensity of hnRNPA2 LC to phase separate, we were only able to perform these analyses at 0 mM NaCl. We found that while tyrosine, arginine, and phosphoserine do not interact with hnRNPA2 LC strongly, addition of phophotyrosine induced LLPS of hnRNPA2 LC and seems to preferentially interact with the arginines of hnRNPA2 LC (Figure S3B-D). These results suggest that, at low salt conditions, arginine to phosphotyrosine interactions are important for hnRNPA2 LC LLPS.

**Figure 3:**
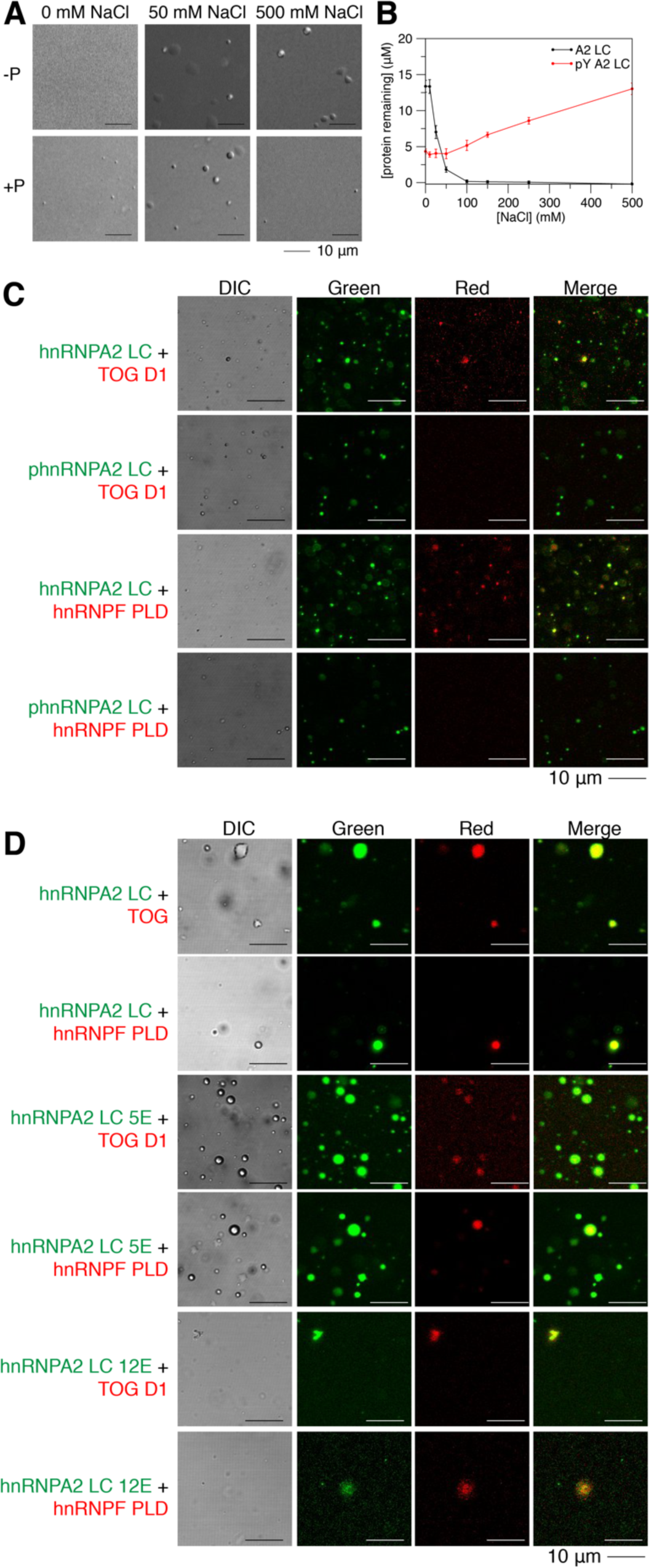
Tyrosine phosphorylation of hnRNPA2 LC alters LLPS and prevents partitioning of hnRNPF PLD or TOG D1. See also Figure S3. (A) While hnRNPA2 LC shows no droplets in low salt conditions and droplets in high salt conditions, tyrosine phosphorylated hnRNPA2 LC shows droplets in all salt concentrations tested, although more droplets are present at low salt conditions. Conditions: 20 µM proteins, 20 mM MES pH 5.5 150 mM urea with 0 mM, 50 mM, or 500 mM NaCl as indicated. Scale bar: 10 µm. (B) Concentration of protein remaining in the supernatant after centrifugation, which is an inverse measure of phase separation, changes with salt concentration. hnRNPA2 LC shows low phase separation (high protein remaining in the supernatant) at low salt conditions and near complete phase separation (no protein remaining in the supernatant) at high salt concentrations. In contrast, tyrosine phosphorylated hnRNPA2 LC shows higher phase separation (less protein remaining in the supernatant) at low salt concentrations than at high salt concentrations. Conditions: 20 µM proteins, 20 mM MES pH 5.5 150 mM urea, salt concentration as indicated, 25°C. (C) Fluorescence micrographs showing that although TOG D1 and hnRNPF PLD partition into hnRNPA2 LC droplets (rows 1 and 3), they are unable to partition into tyrosine phosphorylated hnRNPA2 LC droplets (rows 2 and 4). Conditions: 20 µM proteins (∼1% fluorescently labeled), 20 mM MES pH 5.5, 50 mM NaCl, 150 mM urea. Scale bar: 10 µm. (D) Phosphomimic variants containing 5 or 12 serine to glutamate substitutions (hnRNPA2 LC 5E and 12E, respectively) both allow partitioning of TOG D1 and hnRNPF PLD, indicating that increased negative charge is insufficient to prevent partitioning of TOG D1 and hnRNPF PLD. Conditions: 20 µM proteins (∼1% fluorescently labeled), 20 mM MES pH 5.5, 50 mM NaCl, 150 mM urea. Scale bar: 10 µm.

We next asked if tyrosine phosphorylation of hnRNPA2 LC could alter partitioning of hnRNPF or TOG D1. At 50 mM NaCl both hnRNPA2 LC and pY hnRNPA2 LC undergo LLPS. TOG D1 and hnRNPF PLD each partitioned into hnRNPA2 LC droplets, but interestingly, neither partitioned into pY hnRNPA2 LC (Figure 3C, Figure S3E). Additionally, even hnRNPF FL failed to partition into pY hnRNPA2 LC droplets (Figure S3E). In contrast, the serine phosphomimic constructs 5E and 12E did not prevent partitioning of hnRNPF PLD or TOG D1 (Figure 3D, Figure S3G), suggesting that tyrosine phosphorylation may have a unique role in disrupting hnRNPA2 LC interactions with other hnRNPA2 granule components and that the increased net negative charge arising from phosphorylation is not responsible for the specificity of interactions with hnRNPA2 LC. As arginine was shown to be important for both hnRNPF PLD partitioning and phase separation of phosphotyrosine hnRNPA2 LC, it is possible that hnRNPA2 self-interactions between phosphotyrosine and arginine outcompete the weak interactions with hnRNPF PLD, thus preventing its partitioning. Combined, these results provide residue-specific, mechanistic detail to a previously suggested model in which hnRNPA2 LC tyrosine phosphorylation may play a critical role in regulating LLPS and granule formation/disassociation (Muller et al., 2013; White et al., 2008).

### Tyrosine phosphorylation of hnRNPA2 FL prevents aggregation of disease mutants

Phosphorylation can reduce the aggregation of disease-associated proteins (Monahan et al., 2017). Therefore, we hypothesized that aggregation of hnRNPA2 FL disease variants might be altered by tyrosine phosphorylation. We generated recombinant pY hnRNPA2 FL using the same strategy described for hnRNPA2 LC. After cleavage of a C-terminal maltose binding protein solubility tag, hnRNPA2 FL WT undergoes robust LLPS within 30 minutes, but pY hnRNPA2 FL WT does not phase separate even after two hours (Figure 4A). Consistent with our previous work (Ryan et al., 2018), hnRNPA2 FL D290V forms aggregates after 30 minutes; these aggregates increase in size and number by two hours. In contrast, pY hnRNPA2 FL D290V only formed small droplets after 30 minutes that do not appear to substantially increase in size or form amorphous aggregates in the 2-hour assay period (Figure 4A). Similar to WT, hnRNPA2 FL P298L showed LLPS after 30 minutes but formed aggregates after two hours. Yet, pY hnRNPA2 FL P298L does not undergo LLPS at all, even 2 hours after maltose binding protein cleavage (Figure 4A), like the phosphorylated WT. We conclude that disease variants of hnRNPA2 aggregate over time under these conditions, but increased tyrosine phosphorylation can decrease LLPS and prevent aggregation.

**Figure 4:**
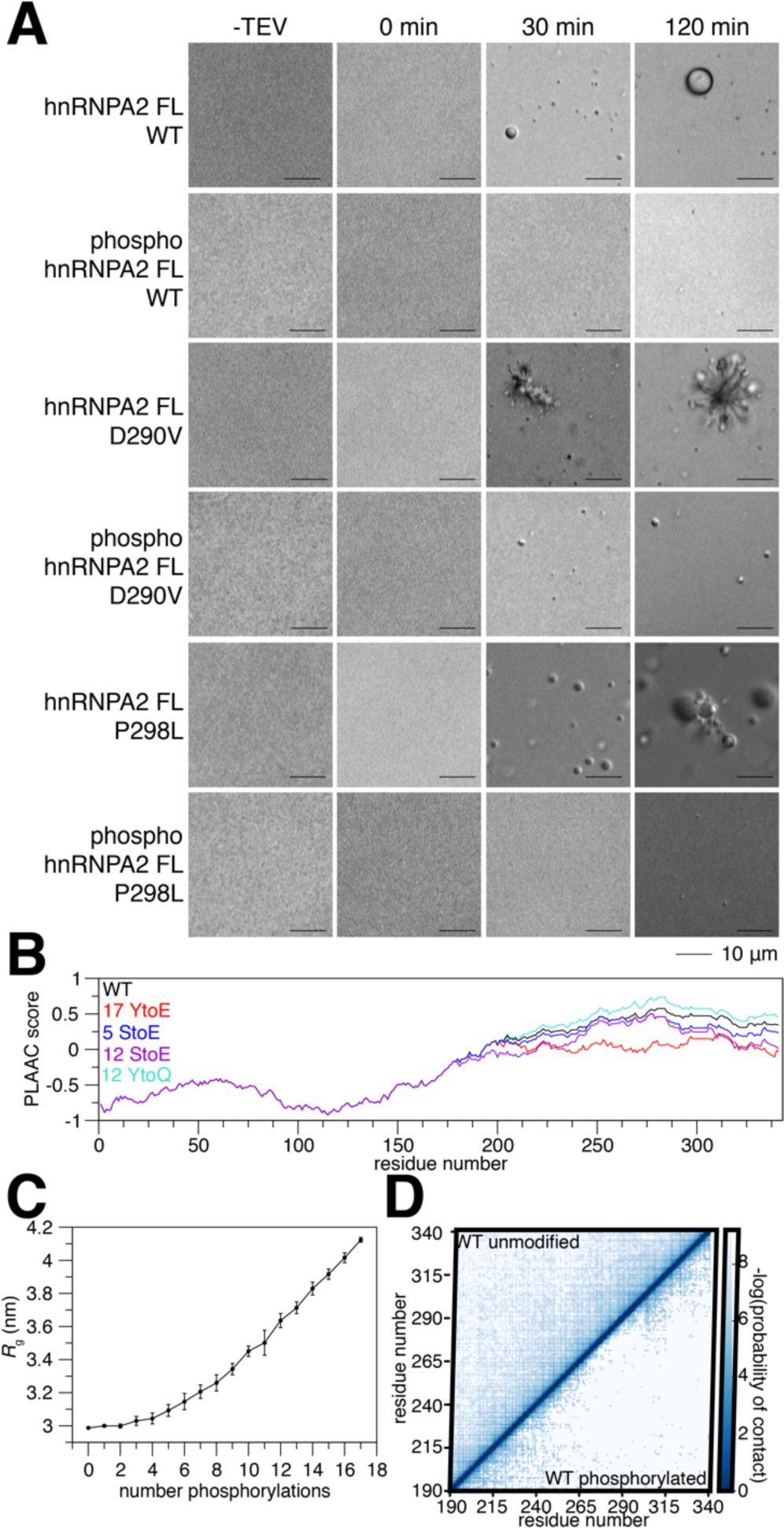
Tyrosine phosphorylation reduces contacts and aggregation of disease mutants. See also Figure S4. (A) After cleavage of a C-terminal maltose binding protein solubility tag, hnRNPA2 FL WT undergoes LLPS and disease mutants D290V and P298L aggregate. In contrast, phosphorylated hnRNPA2 FL WT and P298L do not undergo LLPS or aggregation in the time frame tested. Phosphorylated hnRNPA2 FL D290V can form some structures resembling liquid droplets but does not form amorphous aggregates in the time frame tested. Conditions: 20 µM proteins, 20 mM Tris pH 7.4, 50 mM NaCl. Scale bar: 10 µm (B) PLAAC analysis (Lancaster et al., 2014) of hnRNPA2 (black), changing all tyrosines in hnRNPA2 LC to glutamate (17YtoE, red), 5 serine to glutamate phosphomimic mutations (5StoE, blue), all serines (12StoE, purple), and all tyrosines to glutamine (17YtoQ) shows that hnRNPA2 LC prion-like character reduces slightly with serine phosphomimic mutations, but decreases drastically with tyrosine phosphomimic mutations. (C) As the number of phosphorylation events increases, the radius of gyration (*R*_g_) of hnRNPA2 LC increases in coarse-grained simulations. *R*_g_ uncertainty bars represent S.E.M. from 10 different simulations with the same number of phosphorylated tyrosines randomly placed in different positions. (D) Coarse grained simulations of WT show that compared to the unphosphorylated form, phosphotyrosine hnRNPA2 LC forms fewer contacts with itself.

Given these striking differences in phase separation and aggregation, we sought to test if phosphorylation alters the prion-like character of hnRNPA2. We performed PLAAC analysis (Lancaster et al., 2014) on hnRNPA2 FL and four alternate forms: 1) hnRNPA2 with 5 serine to glutamate mutations in the LC to mimic 5 serine phosphorylations, 2) 12 serine to glutamine mutations to mimic phosphorylation of all serine residues of hnRNPA2, 3) 17 tyrosine to glutamate mutations to attempt to mimic tyrosine phosphorylation of all tyrosines in hnRNPA2 LC, and 4) 17 tyrosine to glutamine mutations to determine the effect of removing the aromatic ring of tyrosine without altering protein charge (because glutamine and glutamate are isosteric, this comparison isolates the importance of the introduction of a negatively charged residue at tyrosine positions in the following analyses). The serine phosphomimics did reduce prion-like character of hnRNPA2 slightly (as compared to non-phosphorylated WT hnRNPA2 FL), but the tyrosine phosphomimic drastically reduced the prion-like character of hnRNPA2 FL, while the tyrosine to glutamine increased prion-like character (Figure 4B). We also performed ZipperDB analysis (Thompson et al., 2006) to see if tyrosine phosphorylation reduces the predicted stability of steric zipper amyloids which are readily formed by the PLD of mutant forms of hnRNPA2 LC (Kim et al., 2013; Ryan et al., 2018). Indeed, tyrosine to glutamate mutations, but not serine to glutamate or tyrosine to glutamine mutations, reduced the predicted favorability of steric zipper formation such that the computed free energy no longer crosses the threshold predicting amyloid formation (Figure S4A). As prion-like character has been suggested to tune the ability to undergo phase separation (Franzmann and Alberti, 2019), this reduction in prion-like character may explain both the altered phase separation and reduced aggregation of the tyrosine phosphorylated hnRNPA2. Finally, we modified an established coarse grain simulation method (Dignon et al., 2018) to incorporate pY residues to evaluate how phosphorylation alters hnRNPA2 LC self-interactions. The disordered monomeric form of hnRNPA2 LC expands as the number of tyrosine phosphorylations increases (Figure 4C), consistent with decreased interactions between the many sites of interaction (Martin et al., 2020). Additionally, hnRNPA2 LC WT, D290V, and P298L all have substantially reducing probability of forming intramolecular contacts when phosphorylated (Figure 4D, S4B-C). These computational assessments further support the model suggesting that tyrosine phosphorylation reduces hnRNPA2 phase separation and aggregation based on our *in vitro* studies.

### Stress alters HRP-1 assembly in *C. elegans* neurons

We next moved to *C. elegans* to assess the impact of tyrosine phosphorylation on hnRNPA2 self-association and toxicity *in vivo*. Although the RRMs of hnRNPA2 homologs are well conserved across animals, disordered domains are usually not well conserved at the amino acid level across species. Therefore, we examined the conservation of hnRNPA2 LC sequence characteristics between humans, vertebrates, and invertebrates. Compared to the average intrinsically disordered region (IDR) (Tompa, 2002) (Figure S5B), human hnRNPA2 LC (Figure S5A) has an unusually high proportion of glycine (46% of residues). We found that the high glycine character was conserved both in vertebrates, with mice, rats, and *Xenopus* all having 46-47% glycine, and in invertebrates, where *Drosophila* Hrb87F and *C. elegans* HRP-1, the orthologs of hnRNAP2, have LC domains with 52-53% glycine (Figure S5A-H). Given this similarity between humans and *C. elegans* in LC domains, we hypothesized that *C. elegans* HRP-1 function is conserved. To test this biochemically, we generated recombinant maltose binding protein-tagged full-length HRP-1 and found that cleavage of the MBP-tag induced HRP-1 phase separation (Figure S5I), as observed previously for human hnRNPA2 FL (Ryan et al., 2018). Additionally, we generated recombinant HRP-1 LC and found this domain was soluble in low salt conditions, like hnRNPA2 LC. At physiological salt, HRP-1 LC formed amorphous aggregates, rather than liquid droplets like hnRNPA2 LC, but both show signs of avid self-assembly (Ryan et al., 2018) (Figure S5J). Given these results, we created *C. elegans* strains expressing a chimeric HRP-1 protein containing most of the human hnRNPA2 LC domain.

To visualize self-association in *C. elegans*, we created transgenic animals expressing HRP-1 tagged with the red fluorescent protein mScarlet (HRP-1mScarlet) (Bindels et al., 2017) inserted between the RRMs and LC (Figure 5A). This construct was expressed in glutamatergic neurons using the *mec-4* promoter and *hrp-1* 3’UTR (Figure 5C). To examine the effect of the D290V disease mutation on self-association, we also replaced the third coding exon of the *C. elegans hrp-1* gene with the corresponding human (*Homo sapiens*, Hs) protein sequence codon optimized for *C. elegans* expression, with the human wild-type sequence (HRP-1HsLC^WT^mScarlet) or with the disease-causing D290V substitution (HRP-1HsLC^D290V^mScarlet). First, we tested whether the *C. elegans* HRP-1mScarlet and chimeric HRP-1HsLC^WT^mScarlet had similar levels of self-association *in vivo* and if exposure to oxidative stress altered self-association in neuronal processes (Figure 5E). As it is unclear if HRP-1 assemblies we observed are stress granules, transport granules, or aggregates, they are referred to herein as “spots”. Without stress, few neuronal processes contained HRP-1mScarlet or HRP-1HsLC^WT^mScarlet spots and there was no difference between transgenes or lines (Figure 5E). After exposure to 22 hours of paraquat-induced oxidative stress, spots increased for both transgenes and all lines. HRP-1mScarlet and HRP-1HsLC^WT^mScarlet were indistinguishable in their response to stress (Figure 5E). We concluded that oxidative stress similarly increases the amount of HRP-1mScarlet and HRP-1HsLC^WT^mScarlet self-association in these neuronal processes and undertook the next studies using only HRP-1HsLC^WT^mScarlet.

**Figure 5:**
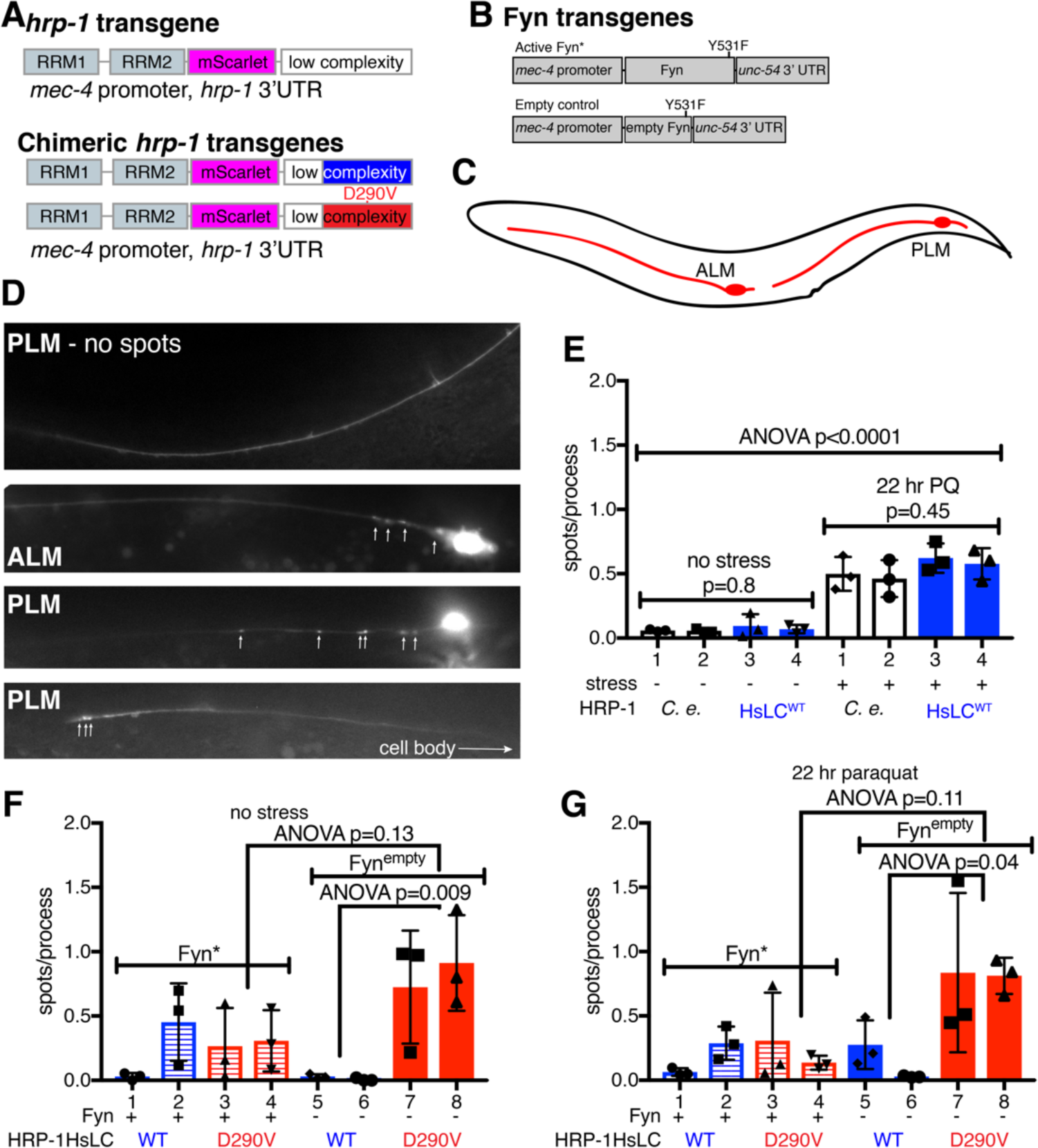
Stress alters HRP-1 assembly *in vivo.* See also Figure S5 and Table S1. (A) mScarlet was added between the RNA recognition motifs (RRMs) and LC of HRP-1 and the third exon was replaced with the corresponding human sequence; disease mutation D290V was introduced. (B) Fyn expression constructs including activated Fyn (Fyn*, Y531F) and empty control Fyn^empty^. (C) Glutamatergic touch neurons ALM and PLM expressing *mec-4* were scored for number of spots in processes. (D) Representative images of *mec-4* neurons with and without HRP-1mScarlet spots. (E) After 22 hours of paraquat induced oxidative stress, both HRP-1mScarlet and HRP-1HsLC^WT^mScarlet assemble more than when unstressed (ANOVA p<0.0001). No significant differences were found within the no stress (p=0.8) stress groups (p=0.45) by ANOVA, indicating that there is no difference in assembly between HRP-1mScarlet and HRP-1HsLC^WT^mScarlet. (F) There is no significant difference in number of spots in HRP-1HsLC^WT^mScarlet lines with or without Fyn, although there seems to be a trend of reduced spots in HRP-1HsLC^D290V^mScarlet lines in animals also expressing Fyn*; this difference is not statistically significant (ANOVA). (G) After exposure to 22 hours paraquat induced oxidative stress there is no significant difference between HRP-1HsLC^WT^mScarlet lines with or without Fyn, although there seems to be a trend of reduced spots in HRP-1HsLC^D290V^mScarlet lines in animals also expressing Fyn*; this difference is not statistically significant (ANOVA). The mean with S.E.M. is reported. Three independent trials for all determinations were performed, with experimenter blinded to genotype for each trial.

To determine if tyrosine phosphorylation alters self-association *in vivo*, we generated transgenes expressing a constitutively active variant of Fyn kinase (Y531F) (Hirose et al., 2003) under the same *mec-4* promoter (Figure 5B). We generated new lines carrying either HRP-1HsLC^WT^mScarlet or HRP-1HsLC^D290V^mScarlet along with either constitutively active Fyn (Fyn*) or an empty control with a frameshift mutation in the Fyn coding region (Fyn^empty^). In the absence of stress, there was no significant difference in spots observed between either HRP-1HsLC^WT^mScarlet or HRP-1HsLC^D290V^mScarlet with or without activated Fyn (Figure 5F). There was not a significant difference after exposure to 22 hours paraquat-induced oxidative stress either (Figure 5G). However, we noted a tendency repeated in both unstressed and stressed conditions for animals expressing both Fyn* and HRP-1HsLC^D290V^mScarlet to have fewer spots than animals expressing HRP-1HsLC^D290V^mScarlet and Fyn^empty^ control. Additionally, HRP-1HsLC^D290V^mScarlet showed significantly more spots than HRP-1HsLC^WT^mScarlet in the absence of Fyn expression (Fyn^empty^) both with and without stress, suggesting that the mutation alters the self-assembly of the chimera in these neurons.

### Chimeric *hrp-1*HsLC^D290V^ causes TDP-43 ortholog-dependent neurodegeneration in *C. elegans* which is rescued by expression of active tyrosine kinase Fyn

Although expression of activated Fyn did not significantly change HRP-1 self-association *in vivo* as monitored by spot formation, we hypothesized that even a slight decrease in D290V association might alter pathological outcomes. To test this, we examined the consequences of manipulating *hrp-1* activity on neurodegeneration. We generated an *hrp-1* genomic rescue construct as before, in which the third coding exon of the *C. elegans hrp-1* gene was replaced with the corresponding human protein sequence (optimized for *C. elegans* expression) resulting in expression of an un-tagged chimeric protein under control of the *hrp-1* promoter and 3’ UTRs (Figure 6A, S6A). We generated additional transgenic lines expressing the disease variant (D290V), lines expressing a truncated form removing the LC domain with a stop codon at the beginning of the 3^rd^ coding exon (ΔLC lines), and control lines with only the plasmid vector and no *hrp-1* DNA (empty). No overt defects were observed in any lines. To assess neurodegeneration, we used dye uptake assays, which provide a sensitive readout of glutamatergic neurodegeneration in other disease models (Baskoylu et al., 2018; Faber et al., 1999). In dye uptake assays, the intact sensory endings of specific glutamatergic sensory neurons take up fluorescent dyes from the environment and backfill cell bodies. Either cell death or process degeneration prevents dye uptake, providing an assessment of neuron integrity. However, even after 22 hours of paraquat-induced oxidative stress, no animals from any line tested had glutamatergic neurodegeneration as assayed by dye update (Figure S6C).

**Figure 6:**
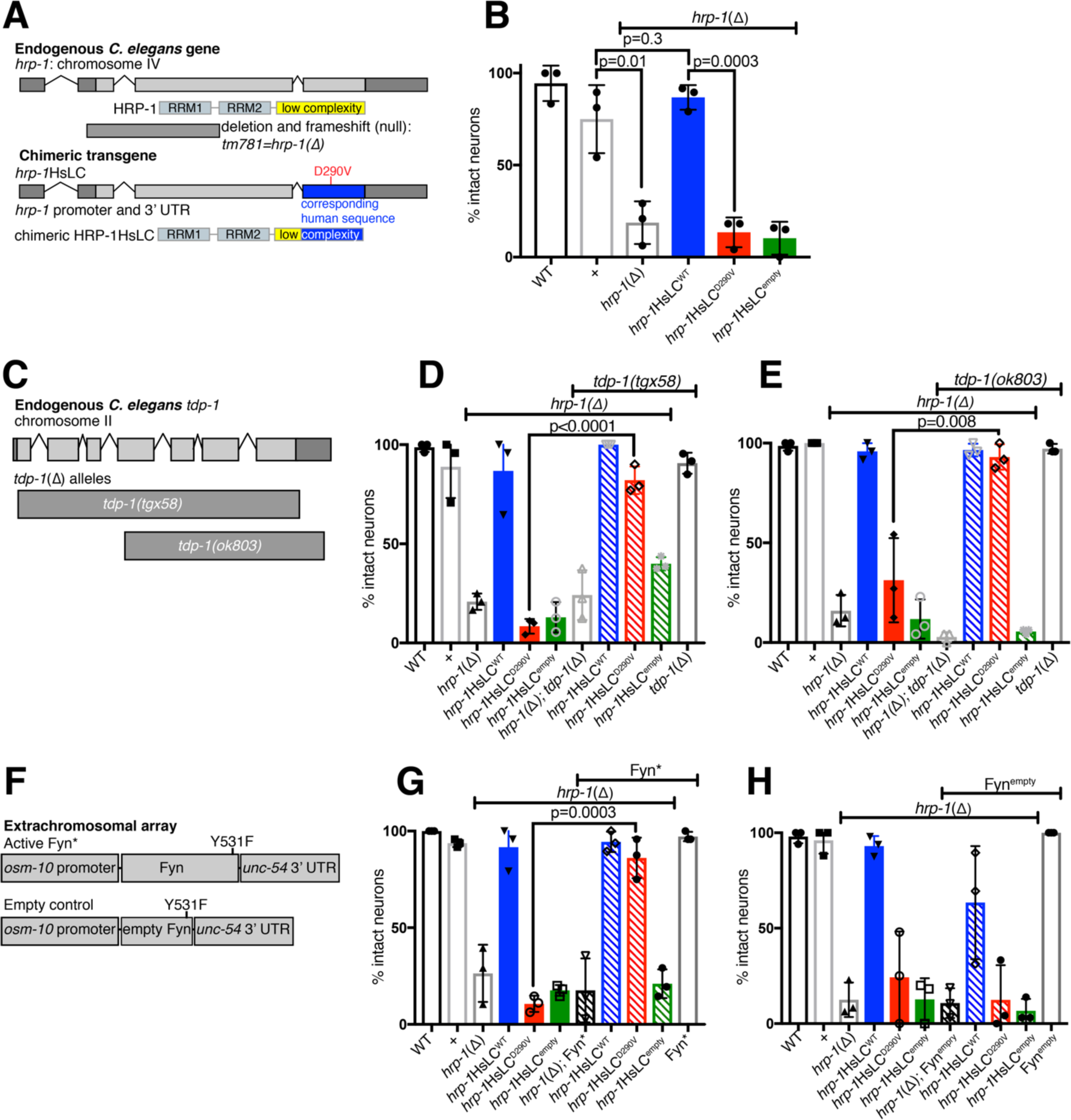
hnRNPA2-D290V causes *tdp-1*-dependent neurodegeneration in *C. elegans*, which is rescued by overexpression of active Fyn kinase. See also Figure S6 and Table S1. (A) Top: schematic depicting *hrp-1* gene and a loss-of-function deletion allele (*tm781*), referred to as *hrp-1*(Δ). Bottom: to create a chimeric protein (HRP-1HsLC), the third coding exon (blue) was replaced with a codon-optimized sequence encoding the majority of the human LC. (B) After 22 hours of paraquat induced oxidative stress, *hrp-1*(Δ) animals showed increased degeneration of tail glutamatergic (phasmid) neurons, based on defective dye uptake. Defective dye uptake indicates either neuron process degeneration or death. This defect is rescued by introduction of *hrp-1*HsLC^WT^, but not *hrp-1*HsLC^D290V^ or *hrp-1*HsLC^empty^. N= 6-12 animals/genotype/trial, significance from two tailed t-test. (C) The *C. elegans* ortholog of TDP-43, *tdp-1*, contains 7 exons. A new deletion allele, *tgx58*, deletes the entire coding region of *tdp-1*. A second deletion allele, *ok803*, removes part of the 4^th^ exon and the remaining three exons. (D) Loss of *C. elegans* TDP-43 ortholog, *tdp-1*, rescues glutamatergic tail/phasmid neurodegeneration caused by *hrp-1*HsLC^D290V^ after 22 hours paraquat induced oxidative stress, but not *hrp-1*(Δ). N=4-12 animals/genotype/trial, significance from two tailed t-test; *tdp-1*Δ=*tdp-1(tgx58).* (E) Loss of *C. elegans* TDP-43 ortholog, *tdp-1*, rescues glutamatergic tail/phasmid neurodegeneration caused by *hrp-1*HsLC^D290V^ after 22 hours paraquat induced oxidative stress, but not *hrp-1*(Δ). N=4-12 animals/genotype/trial, significance from two tailed t-test; *tdp-1*Δ=*tdp-1(ok803).* (F) A constitutively activated Y531F Fyn (Fyn*) construct was expressed in the glutamatergic sensory tail (phasmid) neurons using the *osm-10* promoter. An empty construct removing the beginning of Fyn and causing a frame shift mutation was constructed similarly. (G) Expression of Fyn* in sensory glutamatergic neurons (*osm-10* promoter, *unc-54* 3’UTR) reduces glutamatergic tail/phasmid neuron degeneration in *hrp-1*HsLC^D290V^ animals after 22 hours paraquat induced oxidative stress. N=9-12 animals/genotype/trial, significance from two tailed t-test. (H) Expression of an empty Fyn control (Fyn^empty^) in sensory glutamatergic neurons (*osm-10* promoter, *unc-54* 3’UTR) does not alter glutamatergic phasmid neuron degeneration in *hrp-1*HsLC^D290V^ animals after 22 hours paraquat induced oxidative stress. N=9-12 animals/genotype/trial. The mean with SEM is reported. Three independent trials for all determinations were performed, with experimenter blinded to genotype for each trial. In panels B, D, E, G, H, *hrp-1*HsLC^WT^, *hrp-1*HsLC^D290V^, and *hrp-1*HsLC^empty^ were maintained on a *hrp-1*(Δ)/*tmC25[tmIs1241]* background, but transgenes were tested in animals homozygous for *hrp-1*(Δ). WT is N2, the standard lab strain, while “+” is the N2 *hrp-1* allele maintained over *tmC25[tmIs1241]* and assayed as a homozygote.

**Figure 7:**
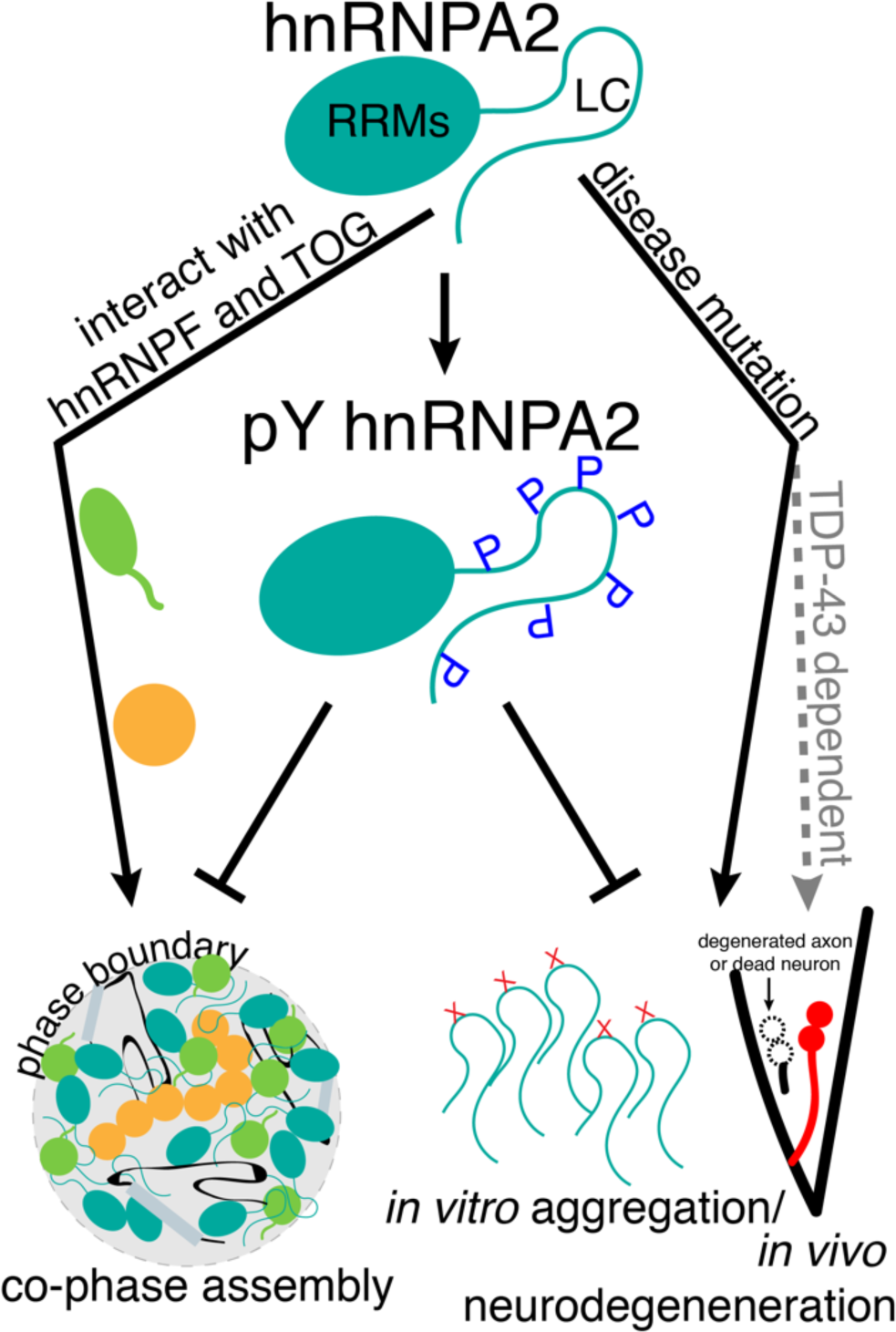
Model: hnRNPA2 interactions and neurodegeneration are altered by tyrosine phosphorylation. hnRNPA2 interacts with transport granule components hnRNPF and ch-TOG in the phase separated state. Disease mutations D290V and P298L induce aggregation of hnRNPA2 *in vitro*, while D290V induces *tdp-1*-dependent neurodegeneration in a *C. elegans* model. hnRNPA2 LC tyrosine phosphorylation alters hnRNPA2 LC LLPS *in vitro*, prevents interaction with hnRNPF and ch-TOG, reduces aggregation *in vitro*, and expression of an activated tyrosine kinase reduces D290V-associated neurodegeneration in the *C. elegans* model.

Realizing that a functional endogenous *C. elegans hrp-1* gene might mitigate the consequences of chimeric HRP-1 transgenes, we crossed the chimeric transgenes onto two available *hrp-1*(Δ) loss-of-function alleles (*tm781* and *ok592*). Knockdown or loss of *hrp-1* causes sterility and partially penetrant lethality (Figure S6B) (Fernandez et al., 2005; Longman et al., 2000; Rual et al., 2004; Sonnichsen et al., 2005); it was important to confirm that the chimeric transgenes rescue this phenotype. Indeed, at least half of the homozygous *hrp-1*(Δ) animals failed to lay eggs by the fourth day of adulthood (Figure S6B), but expression of *hrp-1*HsLC^WT^ restored egg laying (Figure S6B). *hrp-1*(Δ) sterility was also rescued by most of the *hrp-1*HsLC^D290V^ lines (Figure S6B). Neither *hrp-1*HsLC^empty^ or *hrp-1*HsLC^ΔLC^ rescued sterility (Figure S6B), indicating that phenotypic rescue requires an intact LC domain, not simply the RNA-binding regions of HRP-1. Combined, these data show that the glycine-rich LC domain of human hnRNPA2 can substitute *in vivo* for the glycine-rich *C. elegans* HRP-1 LC domain, indicating functional conservation.

Next, we examined glutamatergic neuron degeneration on the *hrp-1* loss-of-function background to determine if *hrp-1*HsLC^D290V^ causes neurodegeneration. Even in the absence of stress, *hrp-1* loss-of-function resulted in partially penetrant dye uptake defects (Figure S6E). Roughly half the tail or head neurons failed to uptake dye (Figure S6E) but the neuron cell bodies were present when scored for expression of *osm-10*p∷GFP (Figure S6F), indicating that the neurons are degenerating but not dying without stress. This dye uptake defect was rescued by introduction of the chimeric *hrp-1*HsLC^WT^ or *hrp-1*HsLC^D290V^ transgenes, but not the empty control *hrp-1*HsLC^empty^ (Figure S6E). Therefore, *hrp-1* is important for neuron survival and D290V does not dramatically impair chimeric HRP-1 function without stress. However, after 22 hours of paraquat-induced oxidative stress, *hrp-1*HsLC^D290V^ animals had increased glutamatergic neurodegeneration in their phasmid neurons (but not amphid neurons), while *hrp-1*HsLC^WT^ animals remained unaffected (Figure 6B, Figure S6D). We conclude that application of oxidative stress reveals neurodegeneration caused by the deleterious impact of the disease-associated D290V mutation on chimeric HRP-1 function.

To reduce the number of alleles tested going forward, we decided to perform further assays on a single *hrp-1*(Δ) allele. In both fertility and glutamatergic neurodegeneration assays, *hrp-1(ok592)* was less severe than *hrp-1(tm781)*, both alone and with *hrp-1*HsLC^D290V^. The *ok592* deletion removes sequences encoding the end of the second RRM domain and the entire LC domain; the first RRM is completely intact. In contrast, *hrp-1(tm781)* is likely a complete loss-of-function (null), as the deletion removes RRM1 coding sequences and frame shifts subsequent translation. Therefore, we used *hrp-1(tm781)* for the remaining studies.

In addition to glutamatergic neurodegeneration, two hallmarks of MSP and associated pathology are cholinergic motor neuron loss and TDP-43 aggregation. To determine the impact of D290V on *C. elegans* motor neuron loss, we used a cholinergic neuron specific GFP transgene to visualize these cells. Both without stress and after 22 hours of paraquat stress, none of the transgenic lines showed cholinergic motor neuron loss compared to control lines (WT or +) (Figure S6G), suggesting that our model does not recapitulate this disease-associated defect. hnRNPA2 and TDP-43 are thought to physically interact to mediate RNA processing (Buratti et al., 2005; D’Ambrogio et al., 2009) and they co-aggregate in patients (Kim et al., 2013) and *in vitro* (Ryan et al., 2018); we hypothesized that loss of *tdp-1*, the *C. elegans* ortholog of TDP-43, might reduce glutamatergic neurodegeneration. Indeed, *tdp-1* loss-of-function did dramatically reduce glutamatergic neurodegeneration in the *hrp-1*HsLC^D290V^ animals but had little or no impact in *hrp-1*(Δ) animals (Figure 6D-E). Two different *tdp-1* loss-of-function alleles (*tgx58* or *ok803*) had the same beneficial impact (Figure 6D-E). As such, *tdp-1* function is required for D290V-associated glutamatergic neurodegeneration in the *C. elegans* models.

As *hrp-1*HsLC^D290V^ animals show neurodegeneration on an *hrp-1* loss-of-function background, we hypothesized that expression of constitutively active Fyn kinase (Fyn*) would alter this neurodegeneration by disrupting aberrant hnRNPA2 D290V interactions. We expressed Fyn* in the glutamatergic neurons affected in *hrp-1*HsLC^D290V^ animals and examined the consequences of Fyn* expression on stress-induced neurodegeneration. We found animals expressing both *hrp-1*HsLC^D290V^ and Fyn* had reduced neurodegeneration when compared to *hrp-1*HsLC^D290V^ animals lacking Fyn expression (Figure 6G). Expression of an empty control (with most of the coding region of Fyn removed, Fyn^empty^) did not alter neurodegeneration in *hrp-1*HsLC^D290V^ animals (Figure 6H). To confirm these results, we made independent transgenes of both Fyn* and Fyn^empty^ and found the same result: active Fyn reduced neurodegeneration in *hrp-1*HsLC^D290V^ animals while Fyn^empty^ did not (Figure S6H). As active Fyn was expressed only in the neurons that degenerate, we can conclude that Fyn has a cell autonomous impact on *hrp-1*HsLC^D290V^-associated neurodegeneration.

## Discussion

Although RNA granules are heterogenous mixtures of proteins and RNAs, much work on LLPS of disease-associated proteins has focused on single proteins (Burke et al., 2015; Conicella et al., 2016; Molliex et al., 2015; Patel et al., 2015; Wang et al., 2018a). Here, we examine the distinct interactions between hnRNPA2 and two other hnRNPA2 transport granule components, hnRNPF and TOG. hnRNPF is readily incorporated into droplets formed by the low complexity domain of hnRNPA2 but not of FUS, suggesting that some specificity of interaction is encoded in distinct prion-like domains. This specificity depends on the arginine residues in hnRNPA2 consistent with the important role for arginine in phase separation due to its unique modes of interaction (Chong et al., 2018; Vernon et al., 2018). In contrast, TOG, an *α*-helix rich globular protein, interacts with hnRNPA2 LC differently than other previously described globular TOG binding partners (Ayaz et al., 2012; Slep and Vale, 2007). These other binding partners, including tubulin, primarily interact with TOG disordered loops (Ayaz et al., 2012) while hnRNPA2 LC binds both the disordered loops and helical faces of TOG D1, which may still permit interaction with tubulin. Of note, the binding affinity for TOG D1 and hnRNPA2 LC is very weak, in contrast to previous reports (Falkenberg et al., 2017). As the previous values were measured with hnRNPA2 FL, it is possible that these measurements were confounded by self-association and LLPS of hnRNPA2, rather than only the stoichiometric heterotypic interaction. Together, these findings show how protein-protein interactions in transport granule form and can give rise to specificity of partitioning despite weak affinities.

Our data suggest that the impact of phosphorylation on LLPS is type and context specific. Serine/threonine phosphorylation in the low complexity domain of FUS reduces aggregation of FUS *in vitro* and *in vivo* (Monahan et al., 2017). Yet, serine/threonine phosphorylation increases LLPS of FMRP (Tsang et al., 2019) and is required for the interaction between FRMP and CAPRIN1 (Kim et al., 2019). We found that although both tyrosine phosphorylation and incorporation of negatively charged amino acids as phosphomimics at serine positions alter LLPS of hnRNPA2 LC, serine phosphomimics do not prevent partitioning of hnRNPF and TOG, while tyrosine phosphorylation does prevent partitioning. This distinction in the ability of related chemical changes to elicit different specific effects suggests that serine and tyrosine phosphorylation may be used to differentially tune interactions between hnRNPA2 and other proteins. Hence, phase separating proteins may encode specificity of partitioning not only by amino acid sequence (see above), but also by post-translational modification of specific residues, significantly expanding the effective amino acid sequence code and providing the ability to dynamic regulate interactions. Of note, tyrosine phosphorylation of ZBP1/IGF2BP1 also leads to local translation of mRNAs carried by this protein (Huttelmaier et al., 2005), suggesting tyrosine phosphorylation may be a common mechanism for initiating translation of transported mRNAs carried in transport granules.

Inhibition of tyrosine kinase pathways and Fyn kinase itself (Imamura et al., 2017; Kaufman et al., 2015; Nygaard, 2018; Nygaard et al., 2014; Smith et al., 2018) are potential therapeutic targets in Alzheimer’s disease. Our data indicate that while tyrosine phosphorylation can reduce aggregation and phase separation *in vitro*, this does not clearly translate to substantial reduction of puncta formation in an *in vivo* model, though the toxic species is not well understood and may not be aggregates or may be smaller than the aggregates probed by fluorescence microscopy (Siddiqi et al., 2019). However, even a modest reduction in self-assembly *in vivo* led to a large reduction in neurodegeneration in an animal model. As such, it is possible that aberrant assembly or aggregation of hnRNPA2 driven by disease mutation is toxic and the presence of these assemblages directly correlates with neurodegeneration. In contrast, it is important to note that activated Fyn may have beneficial functions beyond reducing hnRNPA2 aggregation *in vivo* and, hence, further elucidating the function of Fyn may be of therapeutic interest. These results point to a potential for atomic details of interactions underlying normal granule assembly/disassembly to guide novel strategies to disrupt pathological dysfunction in neurodegenerative disease.

## Acknowledgements

We thank Dr. Vincenzo Venditti for sharing the TROSY-based PRE pulse sequence and Dr. Monica Driscoll for sharing the *mec-4*p∷GFP plasmid. Nemametrix made the *tdp-1(tgx58)* strain in collaboration with the Hart Lab (NIA R43AG061978). Research at Brown University was supported in part by NIGMS R01GM118530 (to NLF), NSF 1845734 (to NLF), and NIH R43AG061978 (to ACH). V.H.R. was supported in part by a Graduate Award from the Robert J. and Nancy D. Carney Institute for Brain Science at Brown University, and Grant F31NS110301 from NINDS, National Institutes of Health. Work at Lehigh University was supported by the U.S. Department of Energy (DOE), Office of Science, Basic Energy Sciences (BES), Division of Material Sciences and Engineering, under Award DESC0013979 (to JM). This research is based in part on data obtained at the Brown University Structural Biology Core Facility supported by the Division of Biology and Medicine, Brown University. Use of the high-performance computing capabilities of the Extreme Science and Engineering Discovery Environment (XSEDE), which is supported by the NSF grant TG-MCB-120014, is gratefully acknowledged in addition to resources of the National Energy Research Scientific Computing Center, a DOE Office of Science User Facility supported by the Office of Science of the U.S. Department of Energy under contract DE-AC02-05CH11231. Some strains were provided by the *Caenorhabditis* Genetics Center (CGC), which is funded by NIH Office of Research Infrastructure Programs (P40 OD010440). Some strains were provided by the National Bioresource Project at the Tokyo Women’s Medical University School of Medicine funded by the Ministry of Education.

## Author Contributions

VHR, ACH, and NLF designed all *in vitro* and *C. elegans* experiments. VHR performed and analyzed all *in vitro* and *C. elegans* experiments, with following exceptions: MTN performed TOG D1 assignment experiments and analyzed data/determined the assignments; CFS performed *osm-10*p∷GFP *C. elegans* experiments. TMP, GLD, and JM designed simulation experiments with input from VHR and NLF. TMP performed and analyzed all simulation experiments with help from GLD. JL aided with *C. elegans* plasmid cloning and strains. VHR, ACH, and NLF wrote the manuscript with input from all authors.

## Conflict of Interests

The authors declare that they have no conflict of interest.

## Methods

### Experimental model and subject details

#### *C. elegans* maintenance

*C. elegans* were maintained at 20°C or 25°C (for *mec-4*p∷*hrp-1*mScarlet extrachromosomal arrays only) under normal growth conditions. Experimenter was blinded to genotype for all trials. All trials were independent biological replicates. See Table 1 for a list of strains. In all figures except Figure S6C, *hrp-1*HsLC^WT^, *hrp-1*HsLC^D290V^, and *hrp-1*HsLC^empty^ were maintained on a *hrp-1*(Δ)*/tmC25[tmIs1241]* background, but transgenes were tested in animals homozygous for *hrp-1*(Δ). *hrp-1*(Δ) was also always maintained over *tmC25[tmIs1241]* but homozygous *hrp-1*(Δ) animals were assayed. In all figures, WT is N2, the standard lab strain, while “+” is the N2 *hrp-1* allele maintained over *tmC25[tmIs1241]* and assayed as a homozygote.

#### Bacterial culture

Uniformly ^15^N, ^13^C, or ^2^H labeled proteins were expressed in M9 in H_2_O or ^2^H_2_O with ^15^N ammonium chloride as the sole nitrogen source or ^13^C-glucose or ^2^H ^13^C-glucose as the sole carbon source as appropriate. Unlabeled proteins were expressed in LB. Cell pellets were harvested from 1 L cultures induced with IPTG at an OD600 of 0.6-1 after 4 hr at 37°C. Tyrosine phosphorylated hnRNPA2 was grown in TKB1 competent cells (Agilent, 200134). 1 L cultures were grown at 18°C overnight to an OD600 of about 1, when there were induced with IPTG for 3 hours at 37°C. Cells were then spun down at 2000 xg for 10 minutes and exchanged into 1x TK media made fresh according to manufacturer’s instructions. Cells were induced in TK media for 2 hours at 37°C before harvesting the cell pellet. Phosphorylated protein yield was low after full purification, so typically 6-12 L of each construct were grown simultaneously to get sufficient yield to freeze purified protein at greater than 1.1 mM. hnRNPA2 LC, FUS LC, hnRNPF PLD, and TOG D1 cell pellets were resuspended in 20 mM NaPi pH 7.4, 300 mM NaCl, 10 mM imidazole, 1 mM DTT while MBP-tagged full length protein pellets were resuspended in 20 mM NaPi pH 7.4, 1 M NaCl, 10 mM imidazole, 1 mM with a Roche Complete EDTA-free protease inhibitor. Resuspended pellets were lysed on an Emulsiflex C3 and the cell lysate cleared by centrifugation (20,000 xg for 60 min at 4°C).

### Method details

#### Recombinant protein

##### Constructs

The following constructs and general purification strategies were used for protein expression in BL21 Star (DE3) *E. coli* cultures (Life Technologies):

- hnRNPA2 LC (190-341), insoluble His-tag purification as described (Ryan et al., 2018) (Addgene ID: 98657)
- hnRNPA2 LC S285C and S329C variants for PRE, insoluble His-tag purification as described (Ryan et al., 2018) (Addgene ID: 98665, 98667 respectively)
- MBP-hnRNPA2 LC, soluble His-tag purification as described (Ryan et al., 2018) (Addgene ID: 98661)
- FUS LC and FUS LC 12E, soluble His-tag purifications as described (Monahan et al., 2017) (Addgene ID: 98653, 98654)
- C-terminal maltose binding protein tagged hnRNPA2 FL WT, D290V, and P298L, soluble His-tag purification (Addgene ID: 139109, 139110, 139111 respectively)
- TOG D1, soluble His-tag purification (Addgene ID: 139112)
- hnRNPF PLD, His-tag purification (Addgene ID: 139113)
- N-terminal maltose binding protein tagged hnRNPF FL and hnRNPF ΔPLD, soluble His-tag purification (Addgene ID: 139114, 139115 respectively)
- HRP-1 LC, insoluble His-tag purification (Addgene ID: 139116)
- C-terminal maltose binding protein tagged HRP-1 FL, soluble His-tag purification (Addgene ID: 139117)
- hnRNPA2 LC no charge, hnRNPA2 LC no charge with R, hnRNPA2 LC RtoK, hnRNPA2 NtoS, hnRNPA2 LC 5E, hnRNPA2 LC 12E, insoluble His-tag purification (Addgene ID: 139118, 139119, 139120, 139121, 139122, 139123 respectively)
- hnRNPF PLD YtoS, hnRNPF PLD StoA, His-tag purification (Addgene ID: 139124, 139125 respectively)
- FUS LC charge, FUS LC charge RtoK, FUS LC with R, insoluble His-tag purification (Addgene ID: 139126, 139127, 139128)

##### Protein purification

hnRNPA2 LC constructs were purified as described (Ryan et al., 2018). FUS constructs were purified as described (Monahan et al., 2017). MTSL labeled hnRNPA2 LC constructs were purified and labeled as described (Ryan et al., 2018). Samples for NMR spectroscopy were produced in M9 minimal media with ^2^H, ^15^N, and ^13^C precursors as appropriate for the experiment.

Briefly, hnRNPA2 LC, variants, and HRP-1 LC were expressed in *E. coli*. Inclusion bodies were resuspended in 8 M urea, 20 mM NaPi pH 7.4, 300 mM NaCl, 10 mM imidazole and cleared by centrifugation at 20,000 xg for 60 min at 4°C. The cleared supernatant was filtered using a 0.2 µm filter and loaded onto a HisTrap 5 mL column. Protein was eluted in a gradient of 10 to 300 mM imidazole over five column volumes. Fractions containing hnRNPA2 LC were pooled, concentrated, and diluted into pH 5.5 MES to a final urea concentration of less than 1 M. Protein was incubated with TEV protease at room temperature overnight. After TEV cleavage, protein was solubilized in 8 M urea and loaded on a HisTrap 5 mL column. Flow through containing cleaved hnRNPA2 LC was collected, concentrated, buffer exchanged into 8 M urea, pH 5.5 MES (pH adjusted with Bis-Tris). Protein was flash frozen at concentrations greater than 1.1 mM.

hnRNPF PLD and variants were purified from *E. coli* lysate by loading cleared soluble lysate onto a HisTrap 5 mL column after filtration by a 0.2 µm filter and then adding the insoluble fraction of the lysate after resuspension in 8 M urea buffer, clearing lysate by centrifugation at 20,000 xg for 60 min at 4°C, and filtering with a 0.2 µm filter. The mixed soluble and insoluble protein was then eluted in a gradient of 10 to 300 mM imidazole (in urea) over 5 column volumes. Collected protein was concentrated and diluted into 20 mM NaPi pH 7.4, 300 mM NaCl to a final imidazole concentration of <50 mM. Protein was incubated with TEV overnight at room temperature. Cleaved protein was filtered then loaded onto a 5 mL HisTrap and flow through containing cleaved protein was collected, concentrated, and buffer exchanged into 20 mM MES pH 5.5 (pH adjusted with BisTris). Protein was flash frozen at concentrations less than 300 µM.

Cleared TOG D1 lysate was loaded onto a HisTrap 5 mL column after filtration. Protein was eluted in a gradient of 10 to 300 mM imidazole over 5 column volumes. Fractions containing TOG D1 were pooled and loaded onto a Superdex 75 sizing column equilibrated in 20 mM NaPi 300 mM NaCl 1 mM DTT. Fractions containing TOG D1 without contaminants were pooled and cleaved by TEV at room temperature overnight. Cleaved protein was filtered and loaded onto a HisTrap 5 mL column and flow through collected, concentrated, and buffer exchanged into 20 mM MES pH 5.5 (pH adjusted in Bis-Tris), and flash frozen at concentrations greater than 1 mM.

Phosphorylated hnRNPA2 LC (and control not phosphorylated protein) was purified from MBP-hnRNPA2 LC. Cleared MBP-hnRNPA2 LC lysate was filtered and loaded onto a HisTrap 5 mL column. Protein was eluted in a gradient of 10 to 300 mM imidazole over five column volumes. Fractions containing MBP-hnRNPA2 LC were pooled and loaded onto a Superdex 200 equilibrated in 20 mM NaPi pH 7.4 300 mM NaCl. Fractions containing MBP-hnRNPA2 LC with minimal degradation were pooled and cleaved with TEV overnight. Protein was solubilized with 8 M urea, filtered, and loaded onto a HisTrap 5 mL. Flow through containing cleaved hnRNPA2 LC was collected, concentrated, and buffer exchanged into 8 M urea 20 mM Bis-Tris pH 5.8.

Protein was loaded onto a monoQ column equilibrated in the same buffer and phosphorylated protein was eluted with a gradient to 1 M NaCl over 7 column volumes. Unphosphorylated protein was collected from the flow through (expressed in BL21 cells due to leaky expression of the tyrosine kinase in TKB1 cells resulting in phosphorylation of hnRNPA2 even in the absence of kinase induction). Collected protein was concentrated, buffer exchanged into 8 M urea pH 5.5 MES (pH adjusted with Bis-Tris) and flash frozen at concentrations greater than 1.1 mM. Cleared maltose binding protein tagged full length protein lysate was filtered and loaded onto a HisTrap 5 mL column. Protein was eluted in a gradient of 10 to 300 mM imidazole over 5 column volumes. Fractions containing MBP-FL were pooled and loaded onto a Superdex 200 sizing column equilibrated in 20 mM NaPi 1 M NaCl pH 7.4. Fractions containing undegraded protein were pooled, concentrated, and flash frozen. Phosphorylated full length hnRNPA2 was further subjected to a monoQ column equilibrated in 20 mM Tris pH 7.5 and eluted with a gradient to 1 M NaCl over 10 column volumes. Phosphorylated protein was collected, concentrated, buffer exchanged into 20 mM NaPi pH 7.4 1 M NaCl and flash frozen.

##### AlexaFluor labeling

Proteins were labeled with NHS-ester AlexaFluors by diluting protein stocks into 20 mM NaPi pH 7.4 300 mM NaCl (1 M NaCl for FL hnRNPA2 and hnRNPF) with 8 M urea for insoluble proteins. AlexaFluor dissolved in DMSO was added at less than 10% total volume. Reactions were incubated for an hour and then free AlexaFluor was removed by desalting with 1 mL Zeba spin desalting columns equilibrated in the appropriate buffer for solubility. Labeled proteins were then concentrated and buffer exchanged into appropriate storage buffers and flash frozen.

##### Phase separation and microscopy

hnRNPA2 LC phase separation assays were performed as described (Ryan et al., 2018). Briefly, protein was diluted from 8 M urea into 20 mM MES pH 5.5 containing the appropriate salt concentration to a final protein concentration of 20 µM and final urea concentration of 150 mM. Samples were left to incubate at room temperature for 10 minutes, then spun down at 17,000 xg for 10 minutes at room temperature. Protein concentration in the supernatant was measured by NanoDrop and calculated using the extinction coefficient of 25330 M^-1^ cm^-1^. Samples were prepared for microscopy by diluting proteins in appropriate conditions to final protein and urea (if applicable) concentrations. DIC images were taken with an Axiovert 200M microscope (Zeiss). Fluorescence microscopy images were taken on an LSM 710 (Zeiss). AlexaFluor-tagged proteins were doped in at 0.2 µL (<1 µM final concentration) to prevent oversaturation of the detector. Snapshots were taken of the red, green, and brightfield channels and merged using ImageJ (NIH).

##### NMR spectroscopy

###### Solution NMR samples

hnRNPA2 LC NMR samples were made by diluting protein from 8 M urea into 20 mM MES pH 5.5 with 10% ^2^H_2_O to a final urea concentration of 150 mM. hnRNPF PLD NMR samples were made by diluting protein from frozen stock (∼300 µM) into 20 mM MES pH 5.5 1 mM DTT with 10% ^2^H_2_O. TOG D1 samples were made by diluting protein from frozen stock into 20 mM MES pH 5.5 1 mM DTT with 10% ^2^H_2_O. Sample concentrations were estimated using the extinction coefficients calculated by ProtParam.

###### Solution NMR experiments

NMR experiments were recorded at 25°C using Bruker Avance III HD NMR spectrometer operating at 850 MHz ^1^H frequency equipped with a Bruker TCI z axis gradient cryogenic probe. Experimental sweep widths, acquisition times, and the number of transients were optimized for the necessary resolution, experiment time, and signal to noise for each experiment type.

###### hnRNPF PLD assignment experiments

Triple resonance assignment experiments were performed on samples of ^13^C/^15^N uniformly labeled hnRNPF PLD (conditions: 20 mM MES pH 5.5 1 mM DTT with 10% ^2^H_2_O). CBCA(CO)NH, HNCACB, HCNO, and HN(CA)CO were recorded with sweep widths 10 ppm (center 4.7) in ^1^H, 20 ppm (center 117) in ^15^N, 6.5 ppm (center 173) in ^13^C for CO experiments and 56 ppm (center 41) in ^13^C for CA/CB experiments using standard Bruker Topspin3.5 pulse programs with default parameter sets (cbcaconhgp3d, hncacbgp3d, hncacogp3d, hncogp3d). Experiments comprised 84-100, 120, 50, and 3072 points in the indirect ^15^N, indirect ^13^Cα/Cβ, indirect ^13^CO, and direct ^1^H dimensions respectively.

###### TOG D1 assignment experiments

Triple resonance assignment experiments were performed on samples of ^2^H/^13^C/^15^N uniformly labeled TOG D1 (conditions: 20 mM MES pH 5.5 1 mM DTT with 10% ^2^H_2_O). HNCA, HN(CO)CA, HNCACB, HN(CO)CACB, HN(CA)CB, HNCO, and HN(CA)CO were recorded with sweep widths 13 ppm (center 4.7) in ^1^H, 25.2 ppm (center 118.65) in ^15^N, 12-13 ppm (center 173) in ^13^C for CO experiments, 60-62 ppm (center 42-44) in ^13^C for CA/CB experiments, and 26-28 ppm (center 53) for CA experiments using standard TROSY-based Bruker Topspin3.5 pulse programs with default parameter sets (trhncagp3d2, trhncocagp3d, trhncacbgp3d, trhncocacbgp3d, trhncacbgp3d, trhncoetgp3d, trhncacogp3d). Experiments comprised 32, 48-64, 96-128, 48, and 1024 points in the indirect ^15^N, indirect ^13^Cα, indirect ^13^Cα/Cβ, indirect ^13^CO, and direct ^1^H dimensions respectively.

###### Paramagnetic relaxation enhancement

Transient intermolecular interactions were probed using paramagnetic relaxation enhancement experiments. The values of the TROSY component of the backbone amide proton transverse relaxation rate constants, ^1^HN *R*_2_, were measured as described (Anthis et al., 2011) using a two dimensional TROSY-based experiment at 850 MHz ^1^H frequency for paramagnetic and diamagnetic samples, with 256 and 3072 total points in the ^15^N indirect and ^1^H direct dimensions, corresponding acquisition times of 44 ms and 139 ms, and sweep widths of 34 ppm and 13 ppm centered around 118 ppm and 4.7 ppm, respectively. Each ^1^H_N_ *R*_2_ experiment comprised six interleaved ^1^H_N_ *R*_2_ relaxation delays in this order: 0 ms, 120 ms, 20 ms, 40 ms, 80 ms, and 10 ms.

### C. elegans

#### Cloning

The following expression constructs were generated:

- pHA#841: *hrp-1*p∷*hrp-1*HsLC^WT^∷*hrp-1* 3’UTR (Addgene ID: 139198)
- pHA#842: *hrp-1*p∷*hrp-1*HsLC^D290V^∷*hrp-1* 3’UTR (Addgene ID: 139199)
- pHA#843: *hrp-1*p∷*hrp-1*HsLC^ΔLC^∷*hrp-1* 3’UTR (Addgene ID: 139200)
- pHA#847: *mec-4*p∷*hrp-1*mScarlet∷*hrp-1* 3’UTR (Addgene ID: 139201)
- pHA#848: *mec-4*p∷*hrp-1*HsLC^WT^mScarlet∷*hrp-1* 3’UTR (Addgene ID: 139202)
- pHA#849: *mec-4*p∷*hrp-1*HsLC^D290V^mScarlet∷*hrp-1* 3’UTR (Addgene ID: 139203)
- pHA#850: *osm-10*p∷FynY531F∷*unc-54* 3’UTR (Addgene ID: 139207)
- pHA#851: *osm-10*p∷Fyn^empty^∷*unc-54* 3’UTR (Addgene ID: 139208)
- pHA#852: *mec-4*p∷FynY531F∷*unc-54* 3’UTR (Addgene ID: 139209
- pHA#853: *mec-4*p∷Fyn^empty^∷*unc-54* 3’UTR (Addgene ID: 139210)

In brief, two fragments containing the *hrp-1* promoter to the beginning of coding exon 3 (4.4 kb) and the 3’UTR (1.2 kb) were amplified from N2 genomic DNA and assembled using the NEBuilder HiFi DNA Assembly kit (E2621S) into pBlueScript with a synthesized DNA fragment containing the human hnRNPA2 LC sequence corresponding to the third coding exon of the *C. elegans* gene, codon optimized for *C. elegans* expression. Blunt end site-directed mutagenesis was used to introduce the D290V mutation and a stop codon at the beginning of the 3^rd^ coding exon for ΔLC. To generate *mec-4p*∷*hrp-1* mScarlet plasmids, mScarlet (Bindels et al., 2017) was amplified from pmScarlet_C1 (Addgene 85042) and assembled into *hrp-1*HsLC^WT^ using NEBuilder HiFi DNA Assembly kit and introducing the D290V mutation by Quickchange. *hrp-1*mScarlet was generated by cloning the *C. elegans* third exon in place of the optimized human sequence using NEBuilder HiFi DNA Assembly kit and then cloning in mScarlet as before. The *hrp-1* promoter was then replaced with the *mec-4* promoter (amplified from *mec-4*p∷GFP, from the Driscoll lab) and assembled into *hrp-1* or *hrp-1*HsLC^WT/D290V^ using NEBuilder HiFi DNA assembly kit. To generate the Fyn expression constructs, Fyn was amplified from mEos2-FYN2-N-10 (Addgene 57380) and assembled into pBlueScript with either a *mec-4* or *osm-10* promoter and the *unc-54* 3’UTR using the NEBuilder HiFi DNA Assembly kit. Blunt end site-directed mutagenesis was performed to introduce the Y531F mutation to generate constitutively active Fyn. To generate Fyn^empty^, the Fyn plasmids were cut with HincII and PflM1, gel extracted, and Klenow filled to generate blunt ends and introduce frame shift mutations before ligating.

#### Strain construction

See Table S1 for detailed genotypes of all strains used or generated for this study. To generate animals expressing mScarlet tagged HRP-1 in MEC-4 touch neurons, we generated extrachromosomal arrays by injecting *pha-1(e2123)* animals with the appropriate *mec-4*p∷*hrp-1*mScarlet∷*hrp-1* 3’UTR construct at 5 ng/µL, *mec-4*p∷GFP at 10 ng/µL, pBX *pha-*1 rescue construct at 100 ng/µL, and 100 ng/µL salmon sperm DNA (to reduce repetitive nature of the array). Animals also expressing a Fyn construct in MEC-4 neurons were generated by injecting *pha-1(e2123)* with the same pools as above with addition of 5 ng/µL of the appropriate *mec-4*p∷Fyn∷*unc-54* 3’UTR construct.

To generate *hrp-1*HsLC animals, we first tried using CRISPR and MosSCI (Zeiser et al., 2011) to generate single copy insertions of chimeric *hrp-1* expressed under its endogenous promoter and 3’ UTR. We could not obtain accurate *hrp-1* homologous recombination events using either method, possibly because of the highly repetitive nature of the endogenous *hrp-1* gene and the repair construct containing the humanized LC. As such, we generated extrachromosomal arrays by injecting into N2 animals with the *hrp-1*HsLC constructs at 25 ng/µL, *elt-2*p∷GFP at 75 ng/µL, and 100 ng/µL salmon sperm DNA (to reduce repetitive nature of the array). The resulting extrachromosomal arrays were randomly integrated into the genome by exposing animals to UV irradiation. Integrated transgenes were backcrossed four times and *hrp-1*(Δ)/balancer was crossed on to generate the final strains for assay.

To generate animals expressing Fyn constructs in *osm-10* neurons, *pha-1(e2123)* animals were injected with 5 ng/µL *osm-10*p∷Fyn∷*unc-54* 3’UTR (either Y531F Fyn* or Fyn^empty^), 2.5 ng/uL PCFJ90 (*myo-2*p∷mCherry), 100 ng/µL pBX1 *pha-1* rescue construct, and 100 ng/µL salmon sperm DNA. Arrays were selected for at 25°C. Resulting arrays express *myo-2*p∷mCherry and were crossed onto integrated *hrp-1*HsLC arrays with *hrp-1*(Δ) *(tm781)*, following the array with *myo-2*p∷mCherry. Animals were grown at 20°C once crosses with *hrp-1*HsLC and *hrp-1*(Δ) were initiated as *pha-1* was not followed.

#### Aggregation

Animals were grown at 25°C because all animals are *pha-1(e2123)* with *pha-1* rescue in the extrachromosomal array. Day 1 adult animals (with or without exposure to 22 hours 2.5 mM paraquat on plates) were picked into 4 µL M9 and immobilized using NemaGel (Nemametrix). Spots of mScarlet tagged HRP-1 (*C. elegans*, HsLC^WT^, or HsLC^D290V^) in the processes of ALM and PLM neurons were counted (63x objective). At least one neuron was counted per animal. Due to the orientation of the animals, often only neurons on one side of the animal could be counted (i.e. 1 ALM and 1 PLM), but sometimes animals were oriented such that 2 neurons of the same type could be counted. There was no difference in number of spots between ALM and PLM. Although they were sometimes visible, PVM and AVM were not scored.

#### Fertility

Twelve L4 animals per genotype (not carrying *the tmC25[tmIs1241]* balancer) were singled to NGM plates seeded with 200 µL OP50 (Day 0). Animals were allowed to grow at 20°C for 4 days. On the fourth day, plates were examined for presence of progeny or eggs. Animals were scored as fertile if eggs or progeny were present and not fertile if no eggs could be found on the plate.

#### Glutamatergic neurodegeneration

Day 1 adult animals were washed off plates with M9 and incubated with DiD (Fisher DilC18(5) D307) in a microfuge tube as in (Perkins et al., 1986). After 1.5 hours, animals were spun down at 10000 rpm for 30 seconds and transferred to a regular NGM plate. After 30 minutes, animals were mounted on 2% (vol/vol) agar pads and immobilized in 30 mg/mL 2-3-butaneione monoxime (BDM, Sigma) in M9 buffer. Fluorescent neuronal cell bodies were visualized and scored for dye uptake under 40x or 63x objectives. There are 4 phasmid neurons per animal, two per side. There are 12 amphid neurons per animal, 6 per side. Neurons were scored as intact if the cell body took up fluorescent dye. Neurons that failed to take up dye could be dead or have degenerated processes. For trials with paraquat stress, animals were pre-exposed to 2.5 mM paraquat for 22 hours on plates.

#### Cholinergic neuronal death assay

A cholinergic (*unc-17p*∷GFP) marker was crossed onto *hrp-1*(Δ) and *hrp-1*HsLC^WT^, *hrp-1*HsLC^D290V^, and *hrp-1*HsLC^empty^ animals. Homozygous *unc-17*p∷GFP marker could not be obtained on the *hrp-1*(Δ);*hrp-1*HsLC background, so animals were carefully picked to ensure they were expressing the marker. Day 1 adult animals were mounted on 2% (vol/vol) agar pads and immobilized with 30 mg/mL 2-3-butaneione monoxime (BDM, Sigma) in M9 buffer.

Fluorescent neurons were visualized and scored at the microscope for cell death based on loss of neuronal GFP under a 63x objective. Animals missing at least two neurons were considered defective. For trails with paraquat stress, animals were exposed to 2.5 mM paraquat for 22 hours on plates.

### Simulations

#### Coarse-grained simulations

Coarse grained simulations on single chains were conducted in cubic boxes with LAMMPS (Plimpton, 1995) software package using our HPS C-α based model (Dignon et al., 2018) coupled to replica exchange molecular dynamics (REMD) (Sugita and Okamoto, 1999) with a temperature list range spanning 150-529K to enhance sampling. HPS treats natural and post-translationally modified residues as single particles. The attractiveness between the particles is scaled based on a common hydrophobicity scale (Kapcha and Rossky, 2014). The hydrophobicity of phosphorylated tyrosine was calculated from ab-initio derived partial charges (Khoury et al., 2013).

#### Coarse-grained simulation analysis

Single chain CG simulations of hnRNPA2 LC were conducted for 1μs for a number of phosphorylated tyrosines ranging from 0 (unmodified) to 17 (fully phosphorylated). 10 additional simulations per number of phosphorylation sites ranging from 1 to 16 were carried to calculate the chain dimension of hnRNPA2 LC wild-type and mutants in intermediate phosphorylation states. Only 10 sequences per phosphorylation site were generated because every possible phosphorylation pattern cannot be sampled. The first 100ns were discarded and the radius of gyration (*R*_g_) was calculated from 5 equal divisions of the equilibrium ensemble at 300K. *R*_g_ uncertainty bars represent SEM from 10 different simulations with the same number of phosphorylated tyrosines randomly placed in different positions. Intramolecular contact maps of hnRNPA2 LC wild-type, D290V and P298L mutants were constructed based on the average number of intramolecular contacts per frame. The distance cut-off for a contact to be formed is defined as two particles within 2σ_ij_ ^⅙^ Å where σ_ij_ is the combined vdW diameter of the two particles (Kim and Hummer, 2008).

#### List of simulations and system sizes presented in this work

*List of simulations and system sizes presented in this work:*
| System | Force field/model | #Atoms/Particles |
| --- | --- | --- |
| hnRNPA2<sup>190-341</sup>WT | CG HPS model | 152 |
| hnRNPA2<sup>190-341</sup>pY | CG HPS model | 152 |
| hnRNPA2<sup>190-341</sup>D290V (unmodified) | CG HPS model | 152 |
| hnRNPA2<sup>190-341</sup>D290V (pY) | CG HPS model | 152 |
| hnRNPA2<sup>190-341</sup>P298L (unmodified) | CG HPS model | 152 |
| hnRNPA2<sup>190-341</sup>P298L (pY) | CG HPS model | 152 |

### Quantification and Statistical analysis

#### *C. elegans* experiments

Data collection and analysis were performed by experimenters blinded to genotype. Quantitative results were analyzed using GraphPad Prism 7. ANOVA was used to determine significance and mScarlet assembly assays. Two tailed t-test was used to determine significance for neurodegeneration assays. A value of *P* < 0.05 was used to establish statistical significance. Error bars in figures represent error or the mean (S.E.M.).

Phase separation

Statistical analysis was performed in Microsoft Office Excel. All data are shown as mean +/-standard deviation.

NMR Spectroscopy

Assignments

Data were processed with nmrPipe (Delaglio et al., 1995) using default linear prediction parameters for either constant time or real-time indirect dimensions and assigned in SPARKY (Lee et al., 2015).

Chemical shift perturbations and intensity ratio

Chemical shifts were calculated by subtracting the chemical shifts of a reference peak from the chemical shift of an experimental peak. Intensity ratios were calculated from the height of each peak and error was propagated from the signal to noise values of each spectrum.

Paramagnetic relaxation enhancement values

Data were processed with nmrPipe (Delaglio et al., 1995), apodized with a cosine squared bell function in the ^1^H dimension and a cosine bell function in the ^15^N dimension. Best-fit relaxation rates were calculated using least-squares optimization of ^1^H/^15^N peak intensities to a single exponential function. PRE rates (Γ_2_) rates were obtained from the difference in ^1^H_N_ *R*_2_ rates for the para-magnetic and diamagnetic samples ^1^H_N_ *R*_2_^para^ - ^1^H_N_ *R*_2_^dia^.

### Data and software availability

NMR chemical shift assignments for hnRNPA2 LC WT were previously deposited (BMRB: 27123) (Ryan et al., 2018) and can be obtained online from the Biological Magnetic Resonance Database (BMRB, http://www.bmrb.wisc.edu/). NMR chemical shift assignments for hnRNPF PLD (BMRB: 50189) and TOG D1 (BMRB deposition in process) and can be obtained online from the BMRB. Plasmids generated herein can be found at Addgene.org. Contact Nemametrix to obtain *tdp-1(tgx58)* animals. Other *C. elegans* strains generated herein can be obtained from the Caenorhabditis Genetics Center (CGC) (https://cgc.umn.edu/strain/search) or by contacting the Hart lab.

## Notes

#### Summary of Updates

Supplemental information (SI figures) added.

## References

Ainger, K., Avossa, D., Morgan, F., Hill, S.J., Barry, C., Barbarese, E., and Carson, J.H. (1993). Transport and localization of exogenous myelin basic protein mRNA microinjected into oligodendrocytes. J Cell Biol 123, 431–441.

Amaya, J., Ryan, V.H., and Fawzi, N.L. (2018). The SH3 domain of Fyn kinase interacts with and induces liquid-liquid phase separation of the low-complexity domain of hnRNPA2. J Biol Chem 293, 19522–19531.

Anthis, N.J., Doucleff, M., and Clore, G.M. (2011). Transient, sparsely populated compact states of apo and calcium-loaded calmodulin probed by paramagnetic relaxation enhancement: interplay of conformational selection and induced fit. J Am Chem Soc 133, 18966–18974.

Ayaz, P., Ye, X., Huddleston, P., Brautigam, C.A., and Rice, L.M. (2012). A TOG:alphabeta-tubulin complex structure reveals conformation-based mechanisms for a microtubule polymerase. Science 337, 857–860.

Banani, S.F., Lee, H.O., Hyman, A.A., and Rosen, M.K. (2017). Biomolecular condensates: organizers of cellular biochemistry. Nat Rev Mol Cell Biol 18, 285–298.

Baskoylu, S.N., Yersak, J., O’Hern, P., Grosser, S., Simon, J., Kim, S., Schuch, K., Dimitriadi, M., Yanagi, K.S., Lins, J., et al. (2018). Single copy/knock-in models of ALS SOD1 in C. elegans suggest loss and gain of function have different contributions to cholinergic and glutamatergic neurodegeneration. PLoS Genet 14, e1007682.

Bindels, D.S., Haarbosch, L., van Weeren, L., Postma, M., Wiese, K.E., Mastop, M., Aumonier, S., Gotthard, G., Royant, A., Hink, M.A., et al. (2017). mScarlet: a bright monomeric red fluorescent protein for cellular imaging. Nat Methods 14, 53–56.

Brumwell, C., Antolik, C., Carson, J.H., and Barbarese, E. (2002). Intracellular trafficking of hnRNP A2 in oligodendrocytes. Exp Cell Res 279, 310–320.

Buratti, E., Brindisi, A., Giombi, M., Tisminetzky, S., Ayala, Y.M., and Baralle, F.E. (2005). TDP-43 binds heterogeneous nuclear ribonucleoprotein A/B through its C-terminal tail: an important region for the inhibition of cystic fibrosis transmembrane conductance regulator exon 9 splicing. J Biol Chem 280, 37572–37584.

Burke, K.A., Janke, A.M., Rhine, C.L., and Fawzi, N.L. (2015). Residue-by-Residue View of In Vitro FUS Granules that Bind the C-Terminal Domain of RNA Polymerase II. Mol Cell 60, 231–241.

Chong, P.A., Vernon, R.M., and Forman-Kay, J.D. (2018). RGG/RG Motif Regions in RNA Binding and Phase Separation. J Mol Biol 430, 4650–4665.

Conicella, A.E., Zerze, G.H., Mittal, J., and Fawzi, N.L. (2016). ALS Mutations Disrupt Phase Separation Mediated by alpha-Helical Structure in the TDP-43 Low-Complexity C-Terminal Domain. Structure 24, 1537–1549.

D’Ambrogio, A., Buratti, E., Stuani, C., Guarnaccia, C., Romano, M., Ayala, Y.M., and Baralle, F.E. (2009). Functional mapping of the interaction between TDP-43 and hnRNP A2 in vivo. Nucleic Acids Res 37, 4116–4126.

Delaglio, F., Grzesiek, S., Vuister, G.W., Zhu, G., Pfeifer, J., and Bax, A. (1995). NMRPipe: a multidimensional spectral processing system based on UNIX pipes. J Biomol NMR 6, 277-293.

Dignon, G.L., Zheng, W., Kim, Y.C., Best, R.B., and Mittal, J. (2018). Sequence determinants of protein phase behavior from a coarse-grained model. PLoS Comput Biol 14, e1005941.

Elvira, G., Wasiak, S., Blandford, V., Tong, X.K., Serrano, A., Fan, X., del Rayo Sanchez-Carbente, M., Servant, F., Bell, A.W., Boismenu, D., et al. (2006). Characterization of an RNA granule from developing brain. Mol Cell Proteomics 5, 635–651.

Faber, P.W., Alter, J.R., MacDonald, M.E., and Hart, A.C. (1999). Polyglutamine-mediated dysfunction and apoptotic death of a Caenorhabditis elegans sensory neuron. Proc Natl Acad Sci U S A 96, 179–184.

Falkenberg, C.V., Carson, J.H., and Blinov, M.L. (2017). Multivalent Molecules as Modulators of RNA Granule Size and Composition. Biophys J 113, 235–245.

Fernandez, A.G., Gunsalus, K.C., Huang, J., Chuang, L.S., Ying, N., Liang, H.L., Tang, C., Schetter, A.J., Zegar, C., Rual, J.F., et al. (2005). New genes with roles in the C. elegans embryo revealed using RNAi of ovary-enriched ORFeome clones. Genome Res 15, 250–259.

Francone, V.P., Maggipinto, M.J., Kosturko, L.D., and Barbarese, E. (2007). The microtubule-associated protein tumor overexpressed gene/cytoskeleton-associated protein 5 is necessary for myelin basic protein expression in oligodendrocytes. J Neurosci 27, 7654–7662.

Franzmann, T.M., and Alberti, S. (2019). Prion-like low-complexity sequences: Key regulators of protein solubility and phase behavior. J Biol Chem 294, 7128–7136.

Gao, Y., Tatavarty, V., Korza, G., Levin, M.K., and Carson, J.H. (2008). Multiplexed dendritic targeting of alpha calcium calmodulin-dependent protein kinase II, neurogranin, and activity-regulated cytoskeleton-associated protein RNAs by the A2 pathway. Mol Biol Cell 19, 2311–2327.

Hirose, T., Koga, M., Ohshima, Y., and Okada, M. (2003). Distinct roles of the Src family kinases, SRC-1 and KIN-22, that are negatively regulated by CSK-1 in C. elegans. FEBS Lett 534, 133–138.

Huttelmaier, S., Zenklusen, D., Lederer, M., Dictenberg, J., Lorenz, M., Meng, X., Bassell, G.J., Condeelis, J., and Singer, R.H. (2005). Spatial regulation of beta-actin translation by Src-dependent phosphorylation of ZBP1. Nature 438, 512–515.

Imamura, K., Izumi, Y., Watanabe, A., Tsukita, K., Woltjen, K., Yamamoto, T., Hotta, A., Kondo, T., Kitaoka, S., Ohta, A., et al. (2017). The Src/c-Abl pathway is a potential therapeutic target in amyotrophic lateral sclerosis. Sci Transl Med 9.

Kapcha, L.H., and Rossky, P.J. (2014). A simple atomic-level hydrophobicity scale reveals protein interfacial structure. J Mol Biol 426, 484–498.

Kaufman, A.C., Salazar, S.V., Haas, L.T., Yang, J., Kostylev, M.A., Jeng, A.T., Robinson, S.A., Gunther, E.C., van Dyck, C.H., Nygaard, H.B., et al. (2015). Fyn inhibition rescues established memory and synapse loss in Alzheimer mice. Ann Neurol 77, 953–971.

Khoury, G.A., Thompson, J.P., Smadbeck, J., Kieslich, C.A., and Floudas, C.A. (2013). Forcefield_PTM: Ab Initio Charge and AMBER Forcefield Parameters for Frequently Occurring Post-Translational Modifications. J Chem Theory Comput 9, 5653–5674.

Kiebler, M.A., and Bassell, G.J. (2006). Neuronal RNA granules: movers and makers. Neuron 51, 685–690.

Kim, H.J., Kim, N.C., Wang, Y.D., Scarborough, E.A., Moore, J., Diaz, Z., MacLea, K.S., Freibaum, B., Li, S., Molliex, A., et al. (2013). Mutations in prion-like domains in hnRNPA2B1 and hnRNPA1 cause multisystem proteinopathy and ALS. Nature 495, 467–473.

Kim, T.H., Tsang, B., Vernon, R.M., Sonenberg, N., Kay, L.E., and Forman-Kay, J.D. (2019). Phospho-dependent phase separation of FMRP and CAPRIN1 recapitulates regulation of translation and deadenylation. Science 365, 825–829.

Kim, Y.C., and Hummer, G. (2008). Coarse-grained models for simulations of multiprotein complexes: application to ubiquitin binding. J Mol Biol 375, 1416–1433.

King, O.D., Gitler, A.D., and Shorter, J. (2012). The tip of the iceberg: RNA-binding proteins with prion-like domains in neurodegenerative disease. Brain Res 1462, 61–80.

Kosturko, L.D., Maggipinto, M.J., D’Sa, C., Carson, J.H., and Barbarese, E. (2005). The microtubule-associated protein tumor overexpressed gene binds to the RNA trafficking protein heterogeneous nuclear ribonucleoprotein A2. Mol Biol Cell 16, 1938–1947.

Kosturko, L.D., Maggipinto, M.J., Korza, G., Lee, J.W., Carson, J.H., and Barbarese, E. (2006). Heterogeneous nuclear ribonucleoprotein (hnRNP) E1 binds to hnRNP A2 and inhibits translation of A2 response element mRNAs. Mol Biol Cell 17, 3521–3533.

Lancaster, A.K., Nutter-Upham, A., Lindquist, S., and King, O.D. (2014). PLAAC: a web and command-line application to identify proteins with prion-like amino acid composition. Bioinformatics 30, 2501–2502.

Laursen, L.S., Chan, C.W., and Ffrench-Constant, C. (2011). Translation of myelin basic protein mRNA in oligodendrocytes is regulated by integrin activation and hnRNP-K. J Cell Biol 192, 797–811.

Lee, W., Tonelli, M., and Markley, J.L. (2015). NMRFAM-SPARKY: enhanced software for biomolecular NMR spectroscopy. Bioinformatics 31, 1325–1327.

Longman, D., Johnstone, I.L., and Caceres, J.F. (2000). Functional characterization of SR and SR-related genes in Caenorhabditis elegans. EMBO J 19, 1625–1637.

Martin, E.W., Holehouse, A.S., Peran, I., Farag, M., Incicco, J.J., Bremer, A., Grace, C.R., Soranno, A., Pappu, R.V., and Mittag, T. (2020). Valence and patterning of aromatic residues determine the phase behavior of prion-like domains. Science 367, 694–699.

Molliex, A., Temirov, J., Lee, J., Coughlin, M., Kanagaraj, A.P., Kim, H.J., Mittag, T., and Taylor, J.P. (2015). Phase separation by low complexity domains promotes stress granule assembly and drives pathological fibrillization. Cell 163, 123–133.

Monahan, Z., Ryan, V.H., Janke, A.M., Burke, K.A., Rhoads, S.N., Zerze, G.H., O’Meally, R., Dignon, G.L., Conicella, A.E., Zheng, W., et al. (2017). Phosphorylation of the FUS low-complexity domain disrupts phase separation, aggregation, and toxicity. EMBO J 36, 2951–2967.

Muller, C., Bauer, N.M., Schafer, I., and White, R. (2013). Making myelin basic protein -from mRNA transport to localized translation. Front Cell Neurosci 7, 169.

Nedelsky, N.B., and Taylor, J.P. (2019). Bridging biophysics and neurology: aberrant phase transitions in neurodegenerative disease. Nat Rev Neurol 15, 272–286.

Nott, T.J., Petsalaki, E., Farber, P., Jervis, D., Fussner, E., Plochowietz, A., Craggs, T.D., Bazett-Jones, D.P., Pawson, T., Forman-Kay, J.D., et al. (2015). Phase transition of a disordered nuage protein generates environmentally responsive membraneless organelles. Mol Cell 57, 936–947.

Nygaard, H.B. (2018). Targeting Fyn Kinase in Alzheimer’s Disease. Biol Psychiatry 83, 369–376.

Nygaard, H.B., van Dyck, C.H., and Strittmatter, S.M. (2014). Fyn kinase inhibition as a novel therapy for Alzheimer’s disease. Alzheimers Res Ther 6, 8.

Patel, A., Lee, H.O., Jawerth, L., Maharana, S., Jahnel, M., Hein, M.Y., Stoynov, S., Mahamid, J., Saha, S., Franzmann, T.M., et al. (2015). A Liquid-to-Solid Phase Transition of the ALS Protein FUS Accelerated by Disease Mutation. Cell 162, 1066–1077.

Perkins, L.A., Hedgecock, E.M., Thomson, J.N., and Culotti, J.G. (1986). Mutant sensory cilia in the nematode Caenorhabditis elegans. Dev Biol 117, 456–487.

Plimpton, S. (1995). Fast Parallel Algorithms for Short-Range Molecular-Dynamics. J Comput Phys 117, 1–19.

Qi, X., Pang, Q., Wang, J., Zhao, Z., Wang, O., Xu, L., Mao, J., Jiang, Y., Li, M., Xing, X., et al. (2017). Familial Early-Onset Paget’s Disease of Bone Associated with a Novel hnRNPA2B1 Mutation. Calcif Tissue Int 101, 159–169.

Raju, C.S., Fukuda, N., Lopez-Iglesias, C., Goritz, C., Visa, N., and Percipalle, P. (2011). In neurons, activity-dependent association of dendritically transported mRNA transcripts with the transacting factor CBF-A is mediated by A2RE/RTS elements. Mol Biol Cell 22, 1864–1877.

Rual, J.F., Ceron, J., Koreth, J., Hao, T., Nicot, A.S., Hirozane-Kishikawa, T., Vandenhaute, J., Orkin, S.H., Hill, D.E., van den Heuvel, S., et al. (2004). Toward improving Caenorhabditis elegans phenome mapping with an ORFeome-based RNAi library. Genome Res 14, 2162–2168.

Ryan, V.H., Dignon, G.L., Zerze, G.H., Chabata, C.V., Silva, R., Conicella, A.E., Amaya, J., Burke, K.A., Mittal, J., and Fawzi, N.L. (2018). Mechanistic View of hnRNPA2 Low-Complexity Domain Structure, Interactions, and Phase Separation Altered by Mutation and Arginine Methylation. Mol Cell 69, 465–479 e467.

Ryan, V.H., and Fawzi, N.L. (2019). Physiological, Pathological, and Targetable Membraneless Organelles in Neurons. Trends Neurosci 42, 693–708.

Shan, J., Munro, T.P., Barbarese, E., Carson, J.H., and Smith, R. (2003). A molecular mechanism for mRNA trafficking in neuronal dendrites. J Neurosci 23, 8859–8866.

Siddiqi, M.K., Malik, S., Majid, N., Alam, P., and Khan, R.H. (2019). Cytotoxic species in amyloid-associated diseases: Oligomers or mature fibrils. Adv Protein Chem Struct Biol 118, 333–369.

Slep, K.C., and Vale, R.D. (2007). Structural basis of microtubule plus end tracking by XMAP215, CLIP-170, and EB1. Mol Cell 27, 976–991.

Smith, L.M., Zhu, R., and Strittmatter, S.M. (2018). Disease-modifying benefit of Fyn blockade persists after washout in mouse Alzheimer’s model. Neuropharmacology 130, 54–61.

Sonnichsen, B., Koski, L.B., Walsh, A., Marschall, P., Neumann, B., Brehm, M., Alleaume, A.M., Artelt, J., Bettencourt, P., Cassin, E., et al. (2005). Full-genome RNAi profiling of early embryogenesis in Caenorhabditis elegans. Nature 434, 462–469.

Sugita, Y., and Okamoto, Y. (1999). Replica-exchange molecular dynamics method for protein folding. Chem Phys Lett 314, 141–151.

Thompson, M.J., Sievers, S.A., Karanicolas, J., Ivanova, M.I., Baker, D., and Eisenberg, D. (2006). The 3D profile method for identifying fibril-forming segments of proteins. Proc Natl Acad Sci U S A 103, 4074–4078.

Tompa, P. (2002). Intrinsically unstructured proteins. Trends Biochem Sci 27, 527–533.

Torvund-Jensen, J., Steengaard, J., Reimer, L., Fihl, L.B., and Laursen, L.S. (2014). Transport and translation of MBP mRNA is regulated differently by distinct hnRNP proteins. J Cell Sci 127, 1550–1564.

Tsang, B., Arsenault, J., Vernon, R.M., Lin, H., Sonenberg, N., Wang, L.Y., Bah, A., and Forman-Kay, J.D. (2019). Phosphoregulated FMRP phase separation models activity-dependent translation through bidirectional control of mRNA granule formation. Proc Natl Acad Sci U S A 116, 4218–4227.

Vernon, R.M., Chong, P.A., Tsang, B., Kim, T.H., Bah, A., Farber, P., Lin, H., and Forman-Kay, J.D. (2018). Pi-Pi contacts are an overlooked protein feature relevant to phase separation. Elife 7.

Wang, A., Conicella, A.E., Schmidt, H.B., Martin, E.W., Rhoads, S.N., Reeb, A.N., Nourse, A., Ramirez Montero, D., Ryan, V.H., Rohatgi, R., et al. (2018a). A single N-terminal phosphomimic disrupts TDP-43 polymerization, phase separation, and RNA splicing. EMBO J 37.

Wang, J., Choi, J.M., Holehouse, A.S., Lee, H.O., Zhang, X., Jahnel, M., Maharana, S., Lemaitre, R., Pozniakovsky, A., Drechsel, D., et al. (2018b). A Molecular Grammar Governing the Driving Forces for Phase Separation of Prion-like RNA Binding Proteins. Cell 174, 688–699 e616.

White, R., Gonsior, C., Bauer, N.M., Kramer-Albers, E.M., Luhmann, H.J., and Trotter, J. (2012). Heterogeneous nuclear ribonucleoprotein (hnRNP) F is a novel component of oligodendroglial RNA transport granules contributing to regulation of myelin basic protein (MBP) synthesis. J Biol Chem 287, 1742–1754.

White, R., Gonsior, C., Kramer-Albers, E.M., Stohr, N., Huttelmaier, S., and Trotter, J. (2008). Activation of oligodendroglial Fyn kinase enhances translation of mRNAs transported in hnRNP A2-dependent RNA granules. J Cell Biol 181, 579–586.

Zeiser, E., Frokjaer-Jensen, C., Jorgensen, E., and Ahringer, J. (2011). MosSCI and gateway compatible plasmid toolkit for constitutive and inducible expression of transgenes in the C. elegans germline. PLoS One 6, e20082.

Zhang, H.L., Eom, T., Oleynikov, Y., Shenoy, S.M., Liebelt, D.A., Dictenberg, J.B., Singer, R.H., and Bassell, G.J. (2001). Neurotrophin-induced transport of a beta-actin mRNP complex increases beta-actin levels and stimulates growth cone motility. Neuron 31, 261–275.

