## Supplementary Figures for "Tyrosine phosphorylation regulates hnRNPA2 granule protein partitioning & reduces neurodegeneration"

### Supplemental figures and table

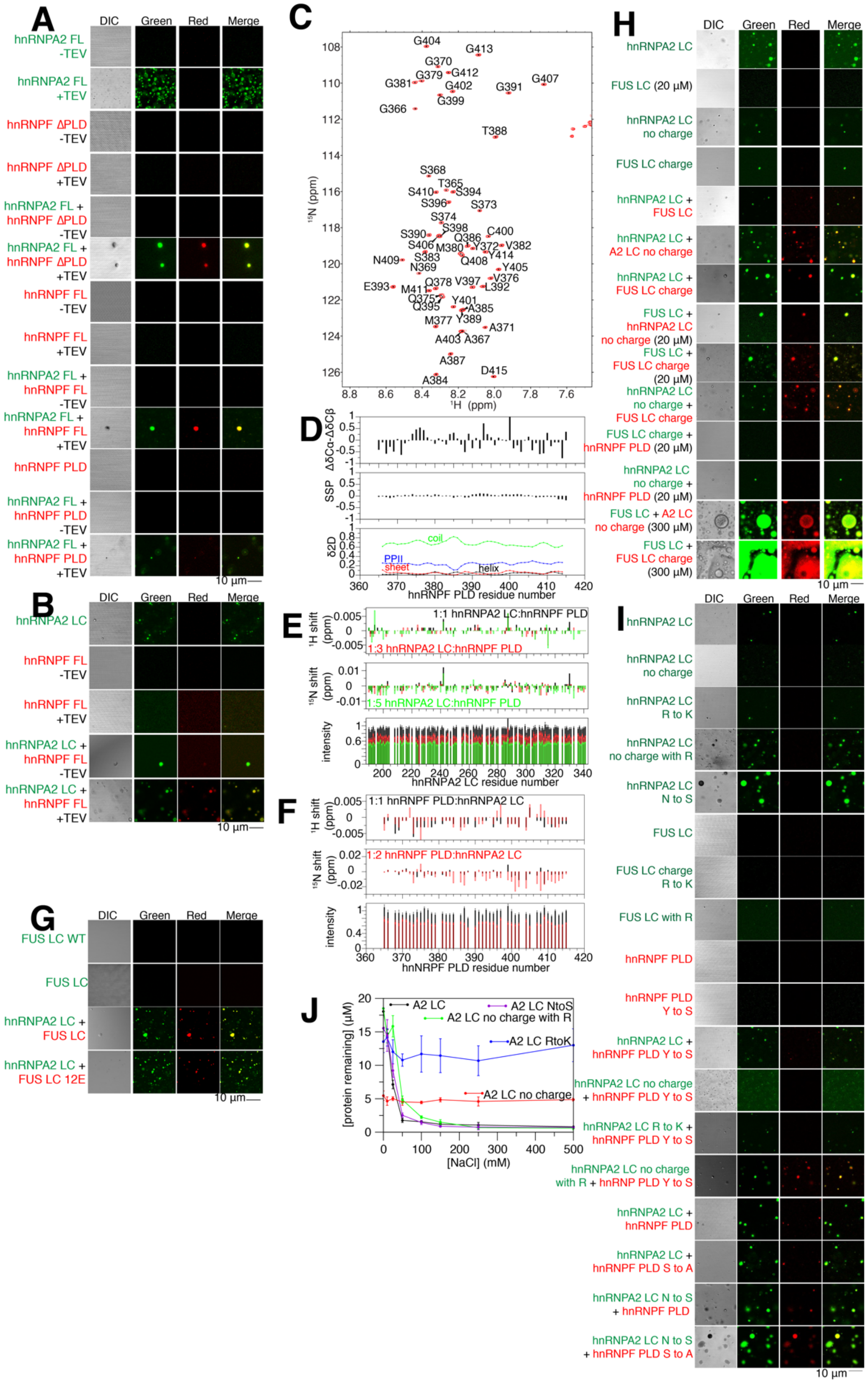

Figure S1: hnRNPF PLD is disordered, induces LLPS of hnRNPA2 LC, and partitioning into hnRNPA2 LC droplets is resistant to hnRNPF PLD mutations. Related to Figure 1.

(A) After cleavage of an N-terminal maltose binding protein solubility tag, hnRNPF FL can partition into hnRNAP2 LC droplets. Scale bar: 10 μm

(B) After cleavage of an N-terminal maltose binding protein solubility tag, hnRNPF FL and hnRNPF ΔPLD do not undergo LLPS. Each one can partition into hnRNPA2 FL droplets. hnRNPA2 FL droplets appear after cleavage of the C-terminal maltose binding protein solubility tag. Scale bar: 10 μm

(C) $^{1}H$-$^{15}N$ HSQC of hnRNPF PLD is consistent with intrinsic disorder.

(D) ΔδCα-ΔδCβ, SSP, and δ2D show hnRNPF PLD is disordered.

(E) Titration of natural isotopic abundance (n.a.) hnRNPF PLD into $^{15}N$ hnRNPA2 LC results in small chemical shifts but decreasing signal intensity with increasing hnRNPF PLD, consistent induction of LLPS.

(F) Titration of natural isotopic abundance (n.a.) hnRNPA2 LC into $^{15}N$ hnRNPF PLD results in small chemical shifts but decreasing signal intensity with increasing hnRNPA2 LC, consistent with induction of LLPS.

(G) FUS LC WT and phosphomimic 12E (Monahan et al., 2017) can partition into hnRNPA2 LC droplets. Scale bar: 10 μm

(H) Brightfield and fluorescence micrographs of control experimental conditions for Figure 3C. Scale bar: 10 μm.

(I) Brightfield and fluorescence micrographs of control experiments for Figure 3G. hnRNPF PLD YtoS and StoA mutations have no effect on partitioning. FUS LC with hnRNPA2 LC-like charge with all arginines changed to lysine is unable to undergo LLPS at 20 μM, as is FUS LC with arginines introduced. Changing the asparagines of hnRNPA2 to serine does not alter partitioning of hnRNPF PLD or hnRNPF PLD StoA, indicating that not all residue substitutions interfere with hnRNPF PLD partitioning into hnRNPA2 LC. Scale bar: 10 μm.

(J) Quantification of phase separation of hnRNPA2 LC constructs used to determine the residue types important for hnRNPF PLD partitioning. hnRNPA2 LC NtoS (purple) has similar phase separation to hnRNPA2 LC. hnRNPA2 LC no charge (red) is consistently phase separated with ~5 μM protein remaining in the supernatant at all salt conditions tested. Adding back arginines to hnRNPA2 LC no charge (hnRNPA2 LC no charge with R, green) brings phase separation as a function of salt to similar levels as hnRNPA2 LC. Changing all the arginine residues to lysine (removing the π-character but maintaining positive charge) also removes the salt dependence of phase separation but has reduced phase separation overall.

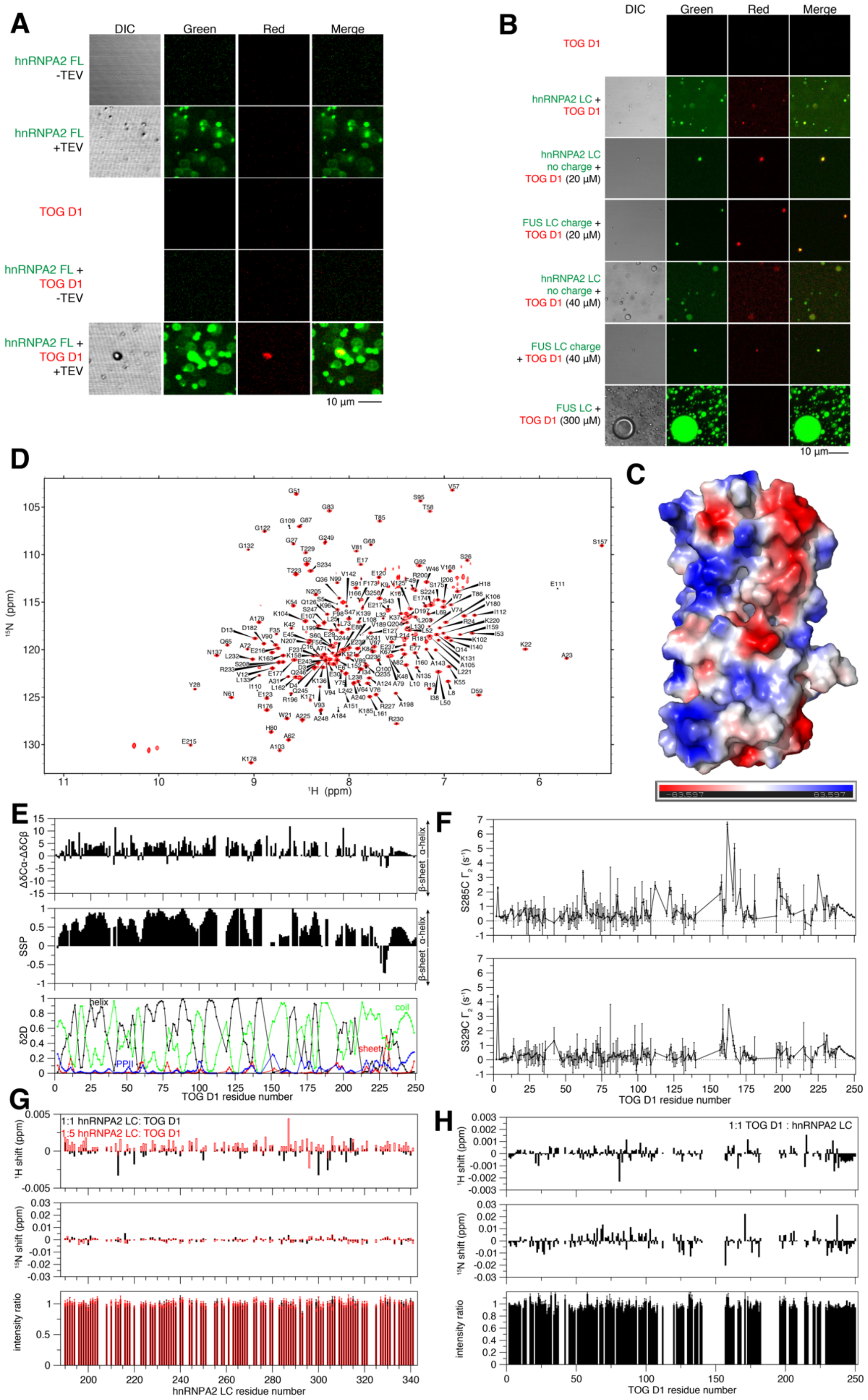


Figure S2: TOG D1 interacts with hnRNPA2 LC weakly. Related to Figure 2.

(A) TOG D1 can partition into hnRNPA2 FL droplets. Scale bar: 10 μm
(B) hnRNPA2 LC no charge and FUS LC charge do not alter partitioning of TOG D1.
(C) TOG D1 homology structure with surface charge indicates that the surface of TOG D1 is highly charged.
(D) $^{1}H$-$^{15}N$ TROSY of $^{2}H$ $^{15}N$ TOG D1 is consistent with a protein with high α-helical content.
(E) ΔδCα-ΔδCβ, SSP, and δ2D indicate that TOG D1 is a globular protein composed mostly of α-helices and disordered linkers.
(F) Quantification of PRE experiments in Figure 2C-D. PREs are overall weaker with hnRNPA2 LC S329C than with hnRNAP2 LC S285C, suggesting that the region around 285 of hnRNPA2 LC interacts with TOG D1 more.
(G-H) Titrations of (G) natural isotopic abundance (n.a.) TOG D1 with $^{15}N$ hnRNPA2 LC and (H) natural isotopic abundance (n.a.) hnRNPA2 LC into $^{2}H$ $^{15}N$ TOG D1 results in small chemical shifts and no change in signal intensity, indicating a very weak interaction.

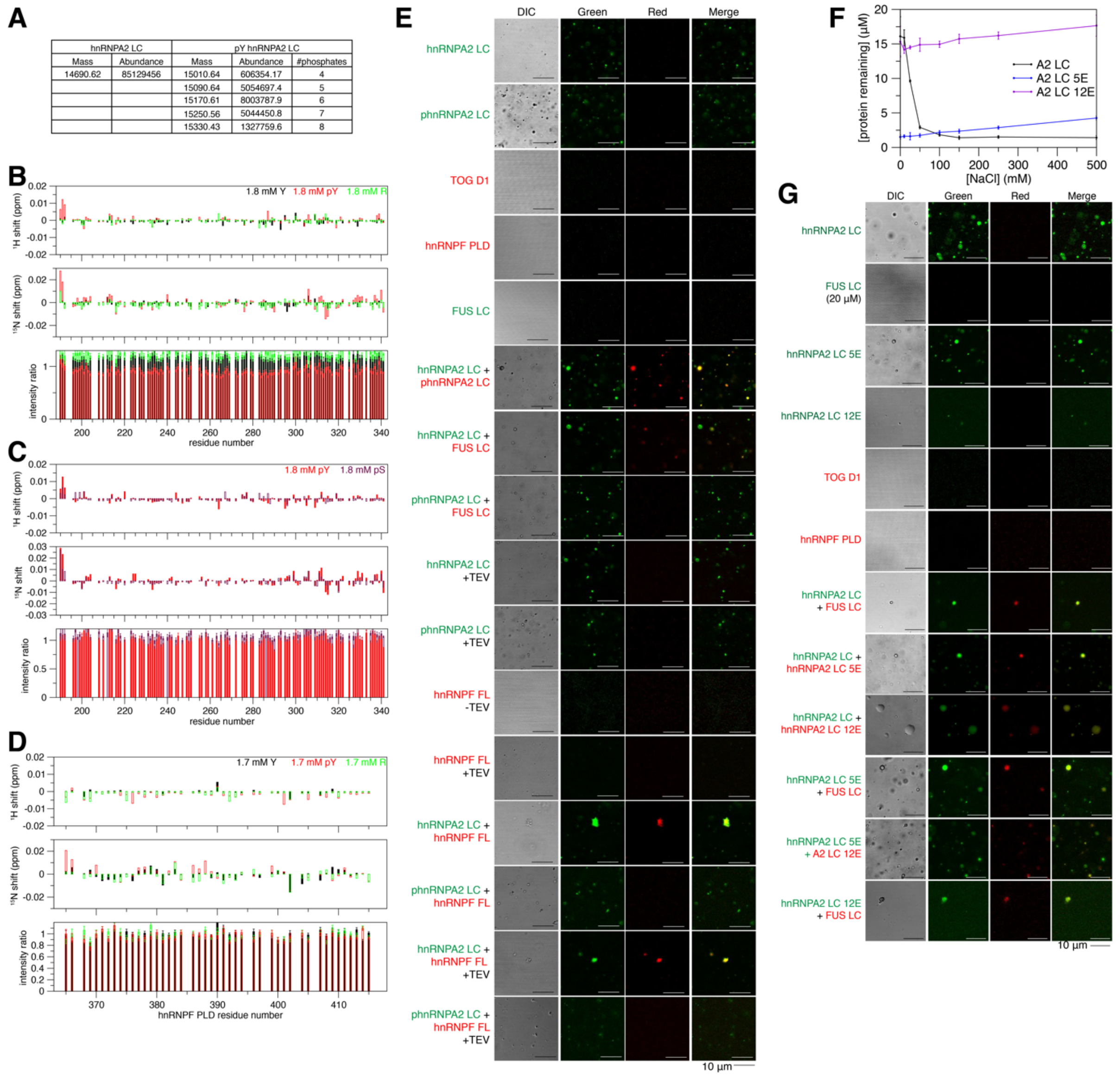


| hnRNPA2 LC | | pY hnRNPA2 LC | | |
|---|---|---|---|---|
| Mass | Abundance | Mass | Abundance | #phosphates |
| 14690.62 | 85129456 | 15010.64 | 606354.17 | 4 |
| | | 15090.64 | 5054697.4 | 5 |
| | | 15170.61 | 8003787.9 | 6 |
| | | 15250.56 | 5044450.8 | 7 |
| | | 15330.43 | 1327759.6 | 8 |

Figure S3: Phosphotyrosine interacts with arginines in hnRNPA2 LC. Related to Figure 3.
(A) Mass spectrometry results showing a range of 4-8 phosphoryl groups have been added to hnRNPA2 LC, with six phosphorylation events being the most abundant.
(B) Titration of free tyrosine, phosphotyrosine, phosphoserine, or arginine with $^{15}$N hnRNPA2 LC. Tyrosine, phosphoserine, and arginine all show small chemical shift perturbations and no intensity changes. In contrast, phosphotyrosine shows larger chemical shifts, particularly in regions of hnRNPA2 LC near arginine residues (indicated) and shows a decrease in signal intensity, indicative of inducing LLPS at these low salt conditions.
(C) Titration of free phosphoserine and phosphotyrosine with $^{15}$N hnRNPA2 LC. Phosphoserine shifts are smaller than those of phosphotyrosine and phosphoserine does not induce LLPS of hnRNPA2 LC.
(D) Titration of tyrosine, phosphotyrosine, and arginine with $^{15}$N hnRNPF PLD. hnRNPF PLD does not phase separate in the presence of any of these amino acids and the chemical shifts are smaller than those of hnRNPA2 LC (see B), but phosphotyrosine induces the largest chemical shifts.

(E) Fluorescence micrographs of control experiments related to Figure 3C and partitioning of hnRNPF FL to hnRNPA2 LC or pY hnRNPA2 LC droplets. hnRNPF FL is also excluded from tyrosine phosphorylated hnRNPA2 LC.
(F) Quantification of phase separation by spinning down phase separated droplets and measuring protein concentration remaining in the supernatant as a function of salt concentration. hnRNPA2 LC with 5 serine to glutamate phosphomimetic mutations (5E, blue) is highly phase separated but phase separation slightly reduces with increasing salt concentration. hnRNPA2 LC with 12 serine to glutamate phosphomimetic mutations (12E, purple) does not phase separate at any salt condition tested.
(G) Fluorescence micrographs of control experiments related to Figure 3D. Serine phosphomimic constructs are able to co-phase separate with WT hnRNPA2 LC and FUS LC. Scale bar: 10 μm.

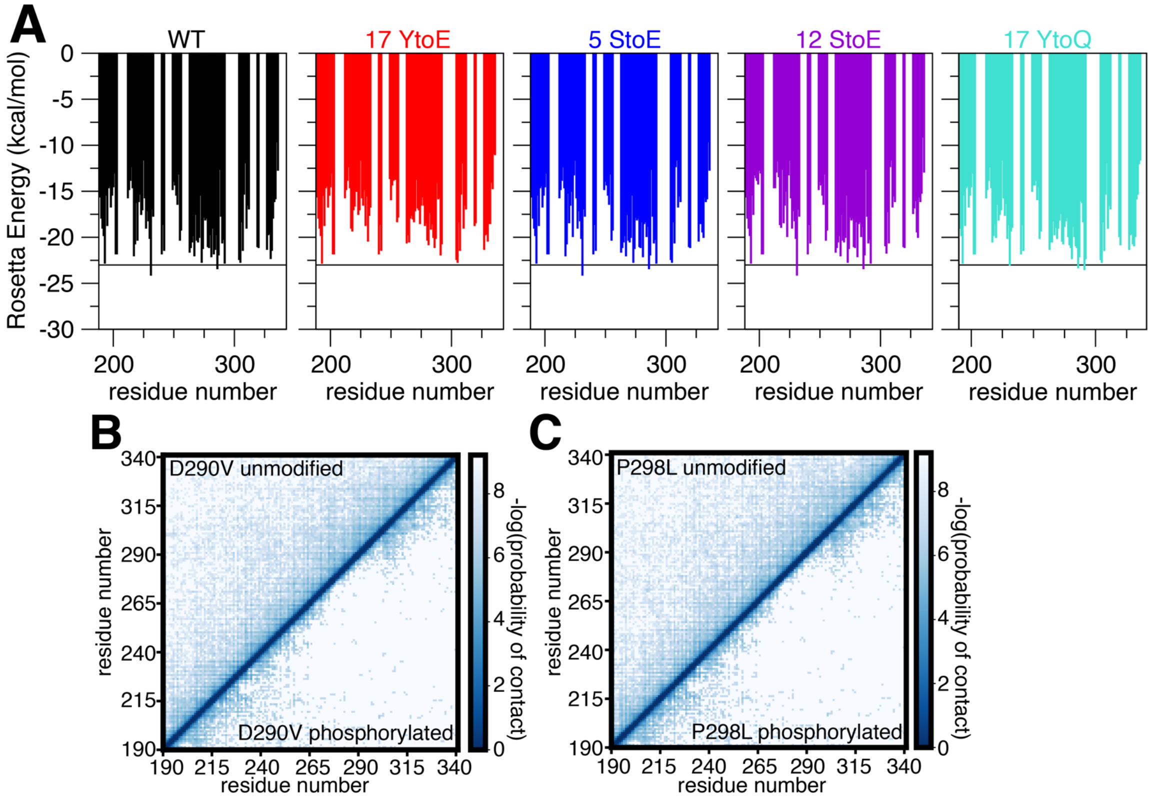


<u>Figure S4: Phosphorylation reduces prion-like character of hnRNPA2.</u> Related to Figure 4.
(A) ZipperDB analysis (Thompson et al., 2006) of the LC domains of the same sequences as in Figure 1D shows decreased Rosetta energy steric zipper forming hexapeptides in the 17YtoE mutant, but not in 5StoE, 12StoE, or 17YtoQ.
(B-C) Coarse grained simulations of D290V (B) and P298L (C) show that compared to the unphosphorylated form, phosphotyrosine hnRNPA2 LC forms fewer contacts with itself.

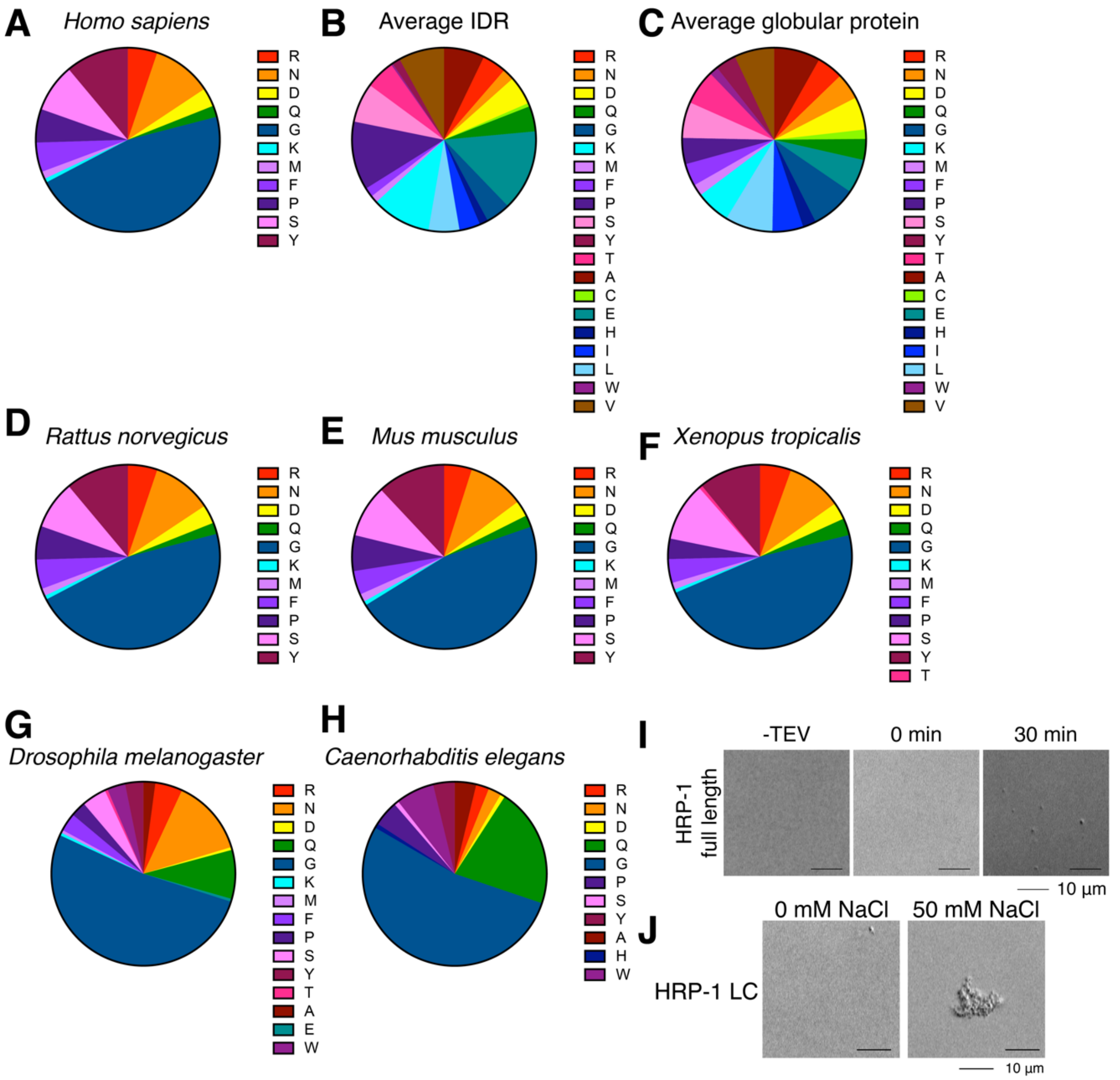


<u>Figure S5: hnRNPA2 LC sequence composition and ability to self-assemble is conserved.</u>
Related to Figure 5.

(A-H) Pie charts of hnRNPA2 LC and ortholog sequence composition. (A) *Homo sapiens* hnRNPA2 LC is predominantly glycine, while (B) the average IDR (Tompa, 2002) and (C) average globular protein (Tompa, 2002) have much greater variation in sequence composition. Vertebrates (D) *Rattus novegicus*, (E) *Mus musculus*, and (F) *Xenopus tropicalis* all show similar hnRNPA2 LC sequence composition to *Homo sapiens*. Invertebrates (G) *Drosophila* and (H) *C. elegans* have a similarly high glycine content in the LC of their hnRNPA2 orthologs.

(I) Recombinant HRP-1 FL is capable of LLPS after cleavage of a maltose binding protein solubility tag. Conditions: 20 µM protein, 20 mM Tris pH 7.4, 50 mM NaCl. Scale bar: 10 µm.

(J) Recombinant HRP-1 LC is soluble at 0 mM NaCl but forms amorphous aggregates in 50 mM NaCl. Conditions: 20 µM protein, 20 mM MES pH 5.5 150 mM urea, salt concentration as indicated. Scale bar: 10 µm

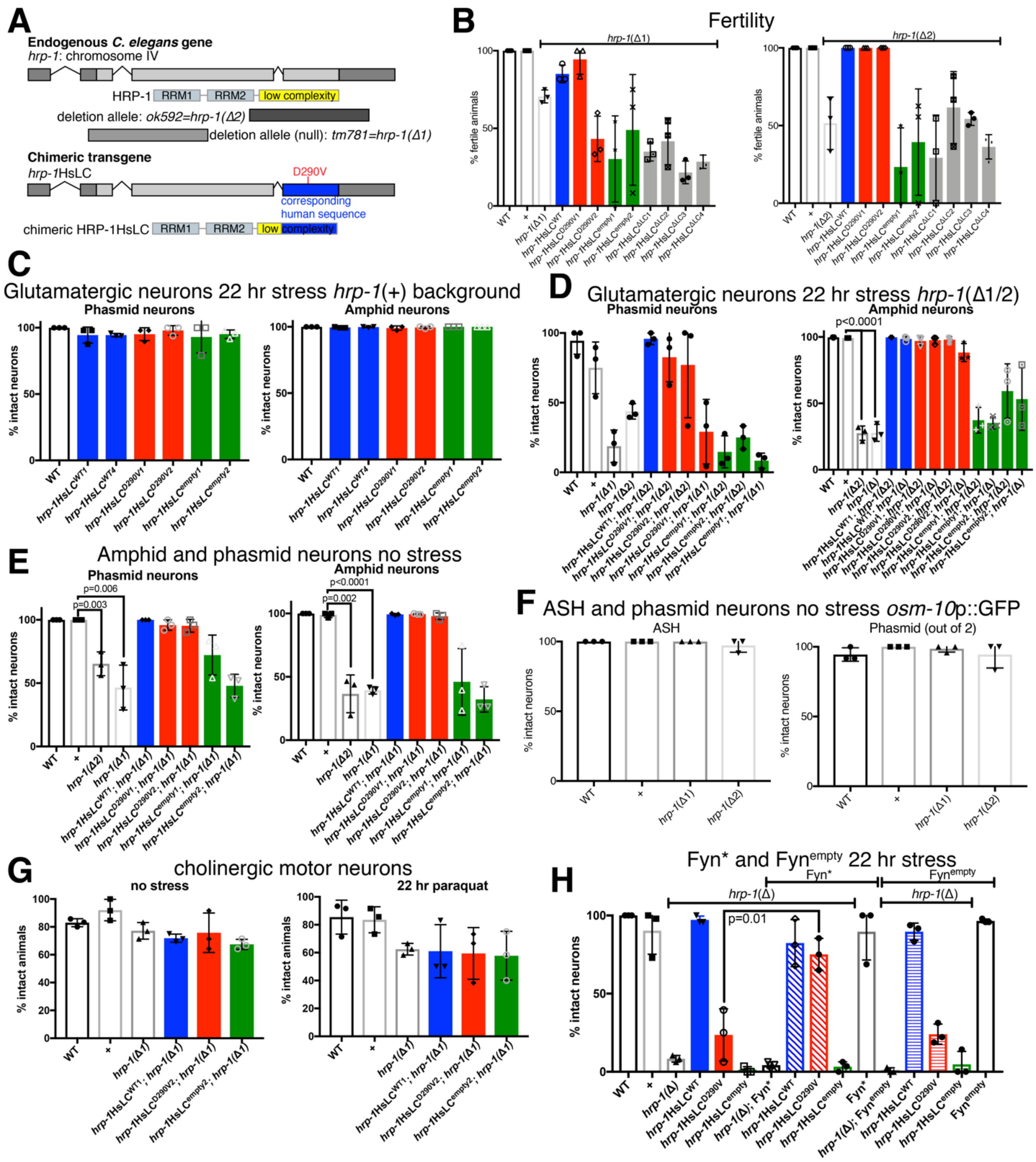


Figure S6: *hrp-1*HsLC$^{D290V}$ animals show glutamatergic neurodegeneration rescued by expression of Fyn kinase. Related to Figure 6, see also Table 1.

(A) Schematic depicting *hrp-1* gene duplicated from Figure 6A with addition of *hrp-1(ok592)* deletion allele.

(B) Animals carrying an *hrp-1(Δ)* allele (*ok592* = 2, *tm781* = 1) do not always lay eggs, thus indicating they have fertility and/or vulval defects. These defects are not rescued by expression of *hrp-1*HsLC$^{\Delta LC}$ or *hrp-1*HsLC$^{empty}$ transgenes. On the *hrp-1*(*ok592*) background, expression of *hrp-1*HsLC$^{WT}$ or *hrp-1*HsLC$^{D290V}$ fully rescued the fertility defect to normal levels. On the *hrp-1*(*tm781*) background, expression of *hrp-1*HsLC$^{WT}$ or *hrp-1*HsLC$^{D290V1}$ transgenes improved fertility, while *hrp-1*HsLC$^{empty}$ or *hrp-1*HsLC$^{D290V2}$ did not, possibly because those animals have multiple protruding vulva (pVul and Muv), which can hamper egg laying. N=12 animals/genotype/trial, 3 trials.

(C) With 22 hours paraquat stress on an *hrp-1(+)* background, *hrp-1*HsLC$^{WT}$, *hrp-1*HsLC$^{D290V}$, or *hrp-1*HsLC$^{empty}$ animals do not have any amphid or phasmid neuron degeneration. N=12 animals/genotype/trial, 3 trials.

(D) Both *hrp-1* deletion alleles (*hrp-1(Δ1) = hrp-1(tm781), hrp-1(Δ2) = hrp-1(ok592)*) have substantial head and tail glutamatergic neuron degeneration after 22 hours of paraquat stress. *hrp-1(tm781)* has more phasmid neuron degeneration than *hrp-1(ok592)* (which likely expresses functional RNA binding domains) both by itself and with expression of *hrp-1*HsLC$^{D290V}$. *hrp-1*HsLC$^{D290V}$ fully rescues the amphid neuron degeneration of both deletion alleles. N=6-12 animals/genotype/trial, 3 trials.

(E) Without exposure to stress, both deletion alleles have milder dye filling defects in both amphid and phasmid neurons, but these defects are fully rescued by expression of *hrp-1*HsLC$^{WT}$ or *hrp-1*HsLC$^{D290V}$. N=10-12 animals/genotype/trial, 3 trials.

(F) Without stress, there is no difference in the number of *osm-10*p::GFP expressing neurons (ASH in the head, 1 phasmid neuron per side in the tail) between WT and either *hrp-1*(Δ) allele, indicating that the neurons are degenerating, not dying. N=12 animals/genotype/trial, 3 trials.

(G) *hrp-1(Δ), hrp-1*HsLC$^{WT}$, *hrp-1*HsLC$^{D290V}$, and *hrp-1*HsLC$^{empty}$ all have similar levels of mild cholinergic neurodegeneration both without stress and after 22 hours of paraquat induced oxidative stress. N=12-16 animals/genotype/trial, 3 trials each no stress and 22 hours paraquat. WT = *hrp-1(+) IV; vsIs48[unc-17p::GFP],* all animals are expressing *vsIs48[unc-17p::GFP]*.

(H) Independent Fyn* and Fyn$^{empty}$ transgenes show the same trend as the initial lines; Fyn* rescues *hrp-1*HsLC$^{D290V}$ neurodegeneration (p=0.01) but not *hrp-1*(Δ), while Fyn$^{empty}$ does not rescue. N=4-12 animals/genotype/trial, 3 trials.

Table S1: Table of *C. elegans* strains used in this study and how they are named in figures.
(see Excel file)
